# The biogeography of evolutionary radiations on oceanic archipelagos

**DOI:** 10.1101/2024.10.07.616413

**Authors:** Baptiste Brée, Thomas J. Matthews, José María Fernández-Palacios, Christian Paroissin, Kostas A. Triantis, Robert J. Whittaker, François Rigal

## Abstract

Evolutionary radiations on oceanic archipelagos (ROAs) have long served as models for understanding evolutionary and ecological processes underlying species diversification. Yet, diversity patterns emerging from ROAs have received relatively little attention from biogeographers, even though characterizing the effect of key geo-environmental factors on island clades species distribution could be important for unraveling diversification dynamics. In this study, we conducted a comparative analysis using island-specific species richness values for approximately one hundred ROAs across major oceanic archipelagos (mostly Hawaii, Canary Islands, Galapagos and Fiji) and taxa (vascular plants, invertebrates and vertebrates). Our aim was to determine whether (1) ROA species richness patterns scale as a function of key geo-environmental factors including island area, geological age, environmental heterogeneity (elevation and topographic complexity) and inter-island isolation, and (2) whether the magnitude of the effects of these factors varies across archipelagos and taxa. Our results identified elevation as a key driver of ROA species richness patterns on islands, supporting existing theoretical and empirical work that highlighted the central role of environmental heterogeneity in driving diversification on oceanic islands. As importantly, we found that the influence of geo-environmental factors varies across archipelagos and taxa, suggesting that unique archipelagic dynamics and biological traits together shape diversification differently. Our findings emphasize the value of applying biogeographical modeling at the resolution of individual radiations to improve our understanding of evolutionary processes on oceanic archipelagos.

## 1. Introduction

Evolutionary radiations on oceanic archipelagos (hereafter ROAs) have served as invaluable models for understanding the eco-evolutionary processes underlying species diversification, due to the high-magnitude geological and environmental changes over evolutionary timescales that characterize volcanic islands (Losos & Ricklefs 2009, Gillespie 2016, Shaw & Gillespie 2016, Whittaker et al. 2023). Here, we define a ROA as a monophyletic clade endemic to an oceanic archipelago, or a group of oceanic archipelagos, that originates from a single colonization event. Since the initial description of the Galápagos finch radiation more than 150 years ago (Darwin 1845, Grant & Grant 2007), many ROAs have been described for multiple taxa and archipelagos, including the iconic Hawaiian Lobeliads (Givnish et al. 2009) and *Drosophila* (O’Grady & DeSalle 2018), and *Naesiotus* land snails of the Galápagos (Parent & Crespi 2006).

Although ROA research has developed considerably in recent decades (e.g. Emerson 2002, Cerca et al. 2023), the diversity patterns emerging from ROAs have attracted surprisingly little attention from island biogeographers. Yet, an evaluation of the geo-environmental factors influencing the island species richness pattern emerging from radiating processes could prove crucial for our understanding of the diversification dynamics taking place in oceanic archipelagos (Whittaker et al. 2008, Economo et al. 2015). In other words, it might be informative to address why, within a given archipelago, some islands have accumulated more species than others during the radiation process. However, despite recent work on oceanic and other island-like systems (e.g. Roell et al. 2021, Wagner et al. 2014, Economo et al. 2015, Lim & Marshall 2017, Tao et al. 2021), evolutionary radiations have rarely been studied from this perspective.

Since the publication of the Equilibrium Theory of Island Biogeography (ETIB) by MacArthur and Wilson (1963, 1967), many strong and predictable relationships between island characteristics and species richness have been identified (Warren et al. 2015). Typically, models applied include island area and isolation as well as environmental heterogeneity and geological age as primary predictors (e.g. Whittaker et al. 2008, 2023, Triantis et al. 2012, Weigelt & Kreft 2013, Barajas-Barbosa et al. 2020). Most of these factors have also been reported as direct or indirect drivers of species diversification on oceanic islands (e.g. Funk & Wagner 1995, Juan et al. 2000, Lim & Marshall 2017, Roell et al. 2021). Therefore, it is reasonable to hypothesise that they will also affect the species richness on islands for a given ROA (hereafter Island S_ROA_). For example, a positive relationship between area and Island S_ROA_ is expected (see Parent & Crespi 2006, Whittaker et al. 2008, Beatty et al. 2017) because larger islands in general offer both more ecological (e.g. diversity of habitats) and geographical (intra-island subdivision of habitats and populations) opportunities for speciation (Losos & Schluter 2000). In addition, the arrangement of many larger oceanic islands within archipelagos also allows for processes of inter-island exchange that further promote radiations (e.g. Grant & Grant 2007).

For isolation, it is therefore primarily the way in which islands are isolated from each other within the archipelago (inter-island isolation) that is expected to determine Island S_ROA_ patterns. Hence, as the ROA spreads throughout an archipelago, species are more likely to colonize the least-isolated islands leading to a negative relationship between within-archipelago inter-island isolation and Island S_ROA_ (Parent & Crespi 2006). Isolation between islands can also enhance or limit diversification by modulating gene flow, promoting or preventing between-island speciation (Hamilton & Rubinoff 1963, Grant & Grant 2007). It is worth noting that propagule exchange may also be influenced by the distribution of island sizes within archipelagos, as larger islands—hosting larger populations—are likely to be more significant sources of colonists than are small islands (Taylor 1987, Weigelt & Kreft. 2013).

Environmental heterogeneity is expected influence rates of key radiation processes as it links speciation rate to two critical island features: (1) the elevation gradient, which serves as a proxy for climatic gradients and habitat diversity, and often promotes speciation by providing ecological opportunities that drive species adaptation (Stuessy et al. 2006), and (2) the topographic complexity that promote within-island allopatry (Whittaker et al. 2008, Kisel & Barraclough 2010). A positive relationship between environmental heterogeneity and Island S_ROA_ is therefore expected. Island geological age might also play a fundamental role in controlling Island S_ROA_. Age has a direct effect on Island S_ROA_ as older islands should (all else being equal) host more species because they have had more time for speciation (Parent & Crespi 2006). However, factors such as area and environmental heterogeneity vary over time following the ontogeny of the islands comprising an archipelago (Whittaker et al. 2008). As they emerge, oceanic islands following typical hotspot dynamics rapidly reach their maximum size and elevation. They then begin to erode and subside, until they become smaller and eventually founder. Topographic complexity will also generally peak during the middle age of the island. As we expect Island S_ROA_ to correlate with elevation and topographic complexity, island age should also have an effect on Island S_ROA_, which should rise and fall throughout the ontogeny of each island, as predicted by the General Dynamic Model of oceanic island biogeography (GDM) (Whittaker et al. 2008, Borregaard et al. 2017).

ROAs are found across a wide array of taxa and archipelagos, raising the question of how the effects of geo-environmental factors on Island S_ROA_ patterns vary across ROAs. A growing body of evidence has substantiated the prevalence of the geo-environmental context in shaping diversification patterns of groups that radiated on oceanic archipelagos (Funk & Wagner 1995, Roman-Palacios & Wiens 2018). It is therefore reasonable to hypothesize that Island S_ROA_ patterns of ROAs belonging to the same archipelago should respond to the distinct combination of island properties in broadly similar ways, thus generating differences in predictors that best explain Island S_ROA_ in different archipelagos. On the other hand, the intrinsic biological characteristics of the ROAs themselves (such as body size and mobility) could also explain why some factors hold greater significance than others. For instance, elevation gradients have been suggested as a primary driver of diversification for several vascular plant ROAs (Itow 1995, Baldwin & Sanderson 1998), for which a positive correlation between Island S_ROA_ and elevation is expected, e.g. as has been found for Hawaiian Lobeliads (Givnish et al. 2009). However, such a correlation might be less prevalent, for example, for a number of arthropod ROAs in Hawaii that appear to have diversified within narrow elevation belts (Hiller et al. 2019).

In this study, we carried out an extensive analysis using a unique dataset that combines Island S_ROA_ values for a hundred ROAs distributed across several oceanic archipelagos and taxa. The aims of our study were twofold, to test whether: (1) Island S_ROA_ scales as a function of different island geo-environmental factors, and (2) the effects of the factors vary depending on the archipelago and taxon to which the ROAs belong. Specifically, we hypothesized that island geo-environmental factors will scale with the Island S_ROA_ of ROAs as outlined in our above predictions (i.e., positive relationships with area, elevation and topographic complexity, negative with inter-island isolation and hump-shaped with geological age). We also hypothesized that, overall, environmental heterogeneity will be the predominant driver of Island S_ROA_ as it is known to trigger diversification across archipelagos and taxonomic groups (Givnish 1997, Losos and Ricklefs 2009, Roell et al. 2021). Lastly, we anticipated that the variation in the contribution of factors would be explained equally by both archipelagos and taxa, as the effect of a factor on Island S_ROA_ stems from complex interactions between the dynamics of the archipelago and the biological characteristics of the ROA itself.

## 2. Material and Methods

### 2.1. Data collection

We have restricted our study to oceanic archipelagos of exclusively volcanic origin that have never been connected to any other landmasses (Triantis et al. 2016). We excluded island systems such as the Philippines that are composed of both volcanic islands and continental fragments (Brown et al. 2013). We then established a list of criteria to determine whether a radiation should be included in our study. First, we selected terrestrial ROAs with a minimum of 10 species. Although arbitrary, this threshold ensures that diversification within the selected ROAs has been sufficiently important for different species to accumulate on several islands within the archipelago, thus providing a minimum of variation in Island S_ROA_ values for the subsequent analyses (see below). For ROAs that diversified across more than one oceanic archipelago, we treated each archipelago independently, only keeping those that harbour a monophyletic clade of at least 10 species (e.g. the *Tarphius* beetle radiation on the archipelagos of Macaronesia (Amorim et al. 2012)). To facilitate comparison between archipelagos, we excluded clades that reach 10 species only at the scale of the meta-archipelago (e.g., South Pacific, Macaronesia). Second, within a given archipelago, the ROA needed to be distributed on at least six islands as any lower threshold is problematic for the regression analyses (Matthews et al. 2023). This threshold also allows for the inclusion of Hawaiian ROAs, as most of them are distributed on the six main islands of the archipelago (i.e. on Kauaʻi, Oʻahu, Molokaʻi, Lāna’i, Maui and Hawai’i; see further details on the island selection procedure for Hawaii in **Appendix 1**). On the other hand, oceanic archipelagos such as Madeira and the Mascarenes, both comprising only 3 islands, were discarded. Third, the ROA should be taxonomically coherent, i.e. all species are from the same genera or from multiple genera that together formed a clade from which the monophyly has been validated by a phylogenetic reconstruction.

To guide our literature search, we first made a list of the major oceanic archipelagos that contain at least six main islands (**Table S1**) and then performed a targeted search of ROA studies with ISI Web of Knowledge and Google Scholar indexing systems (last access December 2022), using archipelago names in combination with “island”, “phylog*”, “radiation” and “endemic” strings. Additionally, we also checked the references list of both the studies we retrieved and some key reviews on the topic (e.g. Shaw & Gillespie 2016, Gillespie et al. 2020, de La Harpe et al. 2017, Florencio et al. 2021, Hembry et al. 2021) to make sure that no relevant ROA had been overlooked. Overall, a total of 105 ROAs distributed across nine oceanic archipelagos were recovered from the literature, Hawaii being by far the most represented of the four with 53 ROAs, followed by the Canary Islands (29), Galápagos (7) and Fiji (6), Society (4), Marquesas (3) and one for Azores, Cabo Verde and Austral Islands (**Figure 1A**). Details on the discarded archipelagos and ROAs are provided in **Table S1** and **Table S2**.

**Figure 1.**
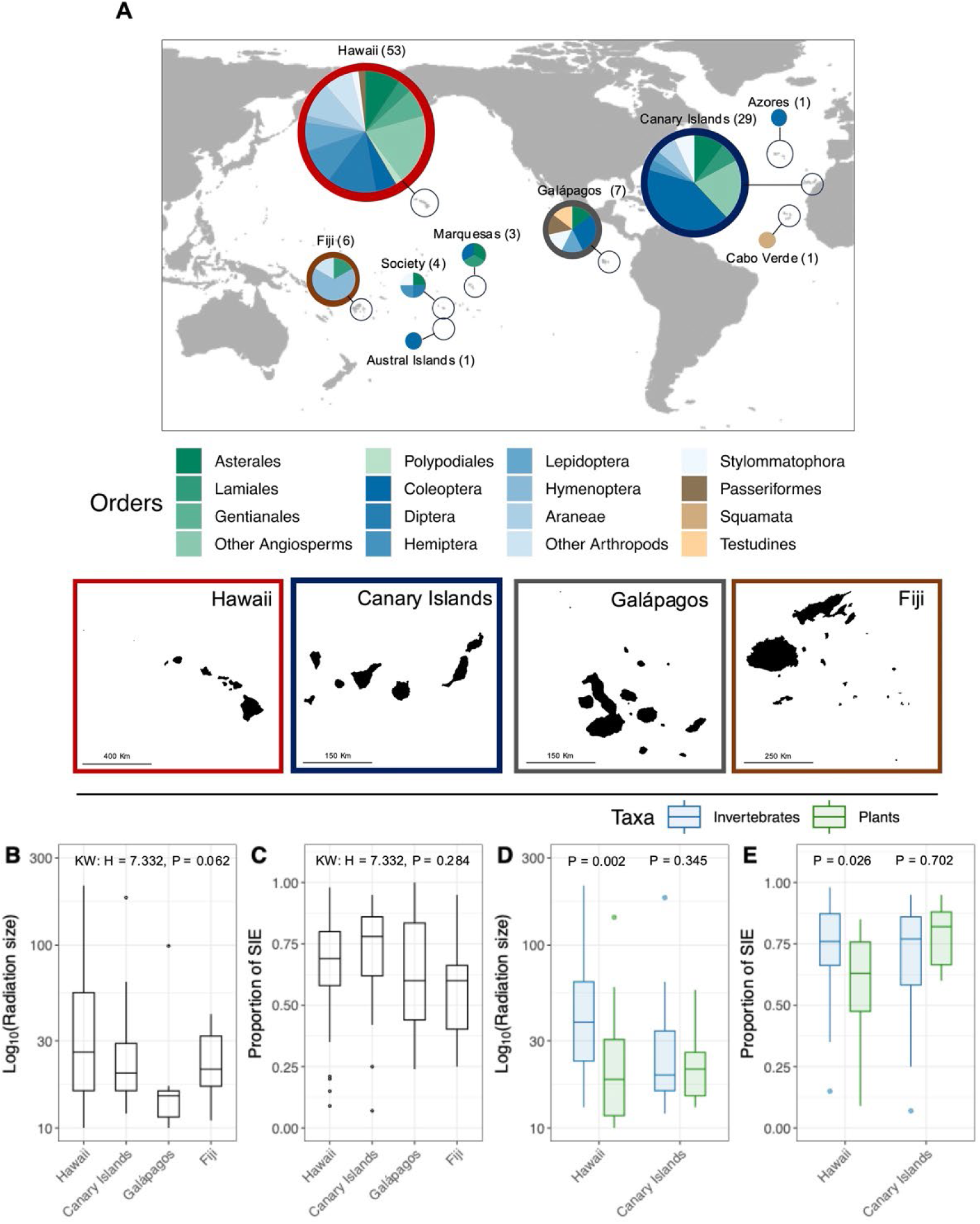
(A) World map indicating the location of the nine archipelagos for which at least one ROA (radiation on oceanic archipelago) fitting our criteria was retrieved. Pie charts showing the proportion of the main orders to which the radiations belong. The four archipelagos analysed in the main text are those with coloured, thick borders and depicted by a map. All together, these four archipelagos represent 90% (95) of all the ROAs retrieved. (B to E). Distribution of (B) the size of the radiation (log_10_-transformed number of species, without subspecies) and (C) the proportion of SIE between archipelagos. The results of a Kruskal-Wallis test of the difference in the size of the radiation and the proportion of SIE between archipelagos is given at the top of each panel. Distribution of (D) the size of the radiation and (E) the proportion of SIE for invertebrates (blue) and plants (green) for both Hawaii and the Canary Islands. The results of Wilcoxon Tests of the difference in the size of the radiation and the proportion of SIE between invertebrates and plants within Hawaii and Canary Islands are given at the top of each panel.

For each selected ROA, we created an occurrence matrix with islands as rows and species as columns for all the species known to belong to the ROA in the focal archipelago, using the most recent taxonomy and sampling published works we found in the literature. When the information was available, we included globally and locally (i.e., species extirpated from individual islands) extinct species. We also created an alternative version of the matrix by including sub-species where they are distinguished, since some authors consider subspecies to be a more appropriate evolutionary unit than species, particularly in the island context (Lasky et al. 2017). The number of species per island was then calculated for each ROA separately to form the Island S_ROA_ vector. For each ROA, we recorded four distinct features: (1) the archipelago it belongs to, (2) taxonomic information (Order, Family, Genus), (3) ROA size (i.e. number of species) and (4) proportion of Single Island Endemic (SIE) species in the ROA. Furthermore, we created a variable named Taxa, where each ROA was classified as either invertebrate, vertebrate or plant. This classification is used here as a coarse proxy for biological characteristics, as previously done in similar contexts (Triantis et al. 2012, Matthews et al. 2016, 2019). Although variability within each taxon may add noise to the analyses, we chose not to use lower ranks in order to maintain sufficient sample sizes. All occurrence matrices are available in **Supplementary Online Data**. The list of the selected ROAs, along with their characteristics and original literature resources is available in **Table S3**.

### 2.2. Island characteristics

Island area and elevation were respectively extracted from Global Administrative Areas (GADM http://www.gadm.org/version1) and the digital elevation model Advanced Land Observing Satellite (ALOS) from the OpenTopography Portal (https://opentopography.org). Island environmental heterogeneity was assessed using (1) the maximum elevation and (2) a measure of topographic complexity, here the variation in the rate of elevational change over the horizontal surface, quantified as the standard deviation of slope (hereafter SDS). Further details are given in **Appendix 1**. Inter-island isolation was calculated using the neighbor index (*NI*) of Kalmar & Currie (2006), which postulates that the importance of an island as a potential source of colonization to the focal island is proportional to its area and to the coast-to-coast distance to the focal island. Including island area in the calculation of inter-island isolation is appropriate, as rates of propagule supply are expected to depend not only on the distance between islands but also on the source size. Larger islands represent more significant sources of colonists, as their larger populations increase the likelihood of dispersal to the focal island. This, in at least some cases, may reduce the probability of extinction on the receiving island (through ‘rescue effects’) and may also sustain gene flow that, in turn, can limit inter-island speciation. Since the perception of inter-island isolation is greatly taxon dependant, *NI* was calculated to give more or less weight to area and distance. To do so, we identified three distinct configurations [i.e. (1) involving only distance, (2) area and distance and (3) area and squared distance] and selected the one that maximized the correlation with Island SROA of each ROA (see further details in **Appendix 1**). Finally, we obtained the geological age of the islands from a literature search. All island characteristics and the associated literature sources are available in **Table S4**. For each archipelago, pairwise correlation between geo-environmental factors was assessed using Pearson’s correlation coefficient (**Figure S1**).

### 2.3. Statistical analyses

All statistical analyses were carried out within the R programming environment (R Development Core Team, 2023). To ensure meaningful between-archipelago comparisons, the analyses below were restricted to the four archipelagos with five or more ROAs (i.e. Hawaii, Canary Islands, Galapagos and Fiji). Collectively, these four archipelagos account for 90% (95) of all the retrieved ROAs. For the five remaining archipelagos (i.e. Society, Marquesas, Austral Islands, Azores, and Cabo Verde), which together represented only 10 ROAs, basic descriptive statistics are given in **Table S5**.

#### 2.3.1 Testing the relationships between geo-environmental factors and Island S_ROA_

We first aimed to investigate univariate relationships between geo-environmental factors and Island S_ROA_ for each ROA separately using the 95 ROAs retrieved from Hawaii, Canary Islands, Galapagos and Fiji. Prior to the analyses, area and elevation were log_10_ transformed to approximate normality. Univariate relationships were assessed using generalized linear models (GLM) using a Poisson distribution or a Negative Binomial distribution (both with a log-link) when overdispersion was detected (Zuur et al. 2009, O’Hara & Kotze 2010). We classified each relationship as (i) being positive or negative and (ii) significant or not. Because a total of 95 GLMs were performed for each factor, we also corrected the *P*-values using the False Discovery Rate method (FDR, Benjamini & Hochberg 1995) to guard against inflation of Type-I errors. Applying such a correction could, however, hamper the detection of important but less strong relationships due to the small number of data points (Borges & Hortal 2009). Therefore, we present the results both with and without the FDR correction. To evaluate the presence of a hump-shaped relationship between age and Island S_ROA_, we performed the GLM with age and its quadratic term (age^2^) and considered the relationship as hump-shaped when it met the three following criteria: (1) the sign of the slope is positive for the linear term and negative for the quadratic term, (2) the two slopes are significant, and (3) the inflection point predicted by the fitted model is present within the range of age values contained in the data (Hortal et al. 2009). To provide an overview of our univariate GLMs, we summed the number of significant positive or negative univariate relationships across archipelagos and taxa for each individual geo-environmental factor. For age + age^2^, we simply counted how many ROAs presented a hump-shaped pattern meeting our three criteria.

#### 2.3.2 Testing whether the magnitude of the effect of geo-environmental factors on Island S_ROA_ varies between archipelagos and taxa

For each geo-environmental factor, we calculated the standardized β coefficient for each relationship established by means of the GLM (above) as a normalized measure of the strength of the effect of each individual factor on Island S_ROA_. The higher or lower the value of the coefficient, the greater the positive or negative effect, respectively. We further computed analyses of variance (ANOVA) for each factor using the standardized β coefficient as the response variable, the archipelago and the taxon to which the ROA belongs to as predictors along with the proportion of SIEs in the ROA and the size of the ROA. For the specific case of age + age^2^, we used instead a binomial GLM with a binary response variable with 1 if S_ROA_ follows a hump-shaped relationship with age and 0 if not. All models were subsequently reduced using backward selection. Initially, we did not include any interaction terms, as the archipelago and taxon variables were both highly unbalanced. Therefore, to test for interactions, we reduced our dataset to only Hawaii and the Canary Islands and to plants and invertebrates in order to obtain a more balanced design. We then reran the ANOVAs as detailed above, testing for interactions between archipelago and taxon only, because preliminary tests for the other potential interactions did not indicate significance.

#### 2.3.3 Quantifying the relative contribution of the geo-environmental factors in explaining Island S_ROA_ variations between archipelagos and taxa

Given that none of the factors is expected to fully account for Island S_ROA_ variation on its own, and that a combination of several (if not all) factors is probably necessary (Roell et al. 2021), we computed Generalized Linear Mixed Models (GLMM) to identify the best set of factors for Island S_ROA_ across ROAs for each of the four archipelagos separately. For Hawaii and the Canary Islands only, we also reran the GLMMs for plant and invertebrate subsets. GLMMs were performed with a Poisson distribution or a negative binomial distribution when overdispersion was detected. The GLMMs were implemented with all geo-environmental factors, including the quadratic term for age to account for potential hump-shaped relationships. We constructed each GLMM by combining all the Island S_ROA_ values belonging to an archipelago (or taxon within an archipelago) in such a way that if, for a given archipelago of six islands, we have recovered 10 ROAs that each radiated over the six islands, the length of the response variable in our GLMM will be 60. ROA identity was added as the random intercept as well as a random slope for all the factors to allow the intercepts and the slopes of the relationships to vary between ROAs. Because the same island appears multiple times in the model, island identity was also included as a separate random effect (intercept only).

The GLMMs were fitted using the R package *glmmTMB* (Brooks et al. 2017), with all factors scaled to zero mean and unit variance. We used a top-down strategy for model selection (Zuur et al. 2009). First, the best random effect structures, with all fixed effects considered, were selected using the *buildglmmTMB* function in the R package *buildmer* (Voeten 2023). Next, we used the *dredge* function in the *MuMIn* package (Bartoń 2023) to fit all possible combinations of fixed effects and ranked the models using their value of AIC corrected for sample size (henceforth AIC_c_). We considered all models with ΔAIC_c_ < 2 (the difference between each model’s AIC_c_ and the lowest AIC_c_) as best models. Note that the term age^2^ was included in a model only where age was included. For each best model we used the *squaredGLMM* function to compute the conditional R^2^ (R^2^c) (Nakagawa & Schielzeth 2013), partitioned into the marginal R^2^ (R^2^m), representing the variance explained by fixed factors and the random R^2^ (R^2^r = R^2^c - R^2^m), representing the variance explained by the random factors. Finally, we derived a measure of variable importance by summing up AIC_c_-weighted standardized regression coefficients for all best models and by scaling them to sum to one for comparison across archipelagos and taxa (Weigelt et al. 2016).

## 3. Results

None of the analyses carried out using subspecies data showed any specific differences from our main results. We therefore present the former only in the Supporting Information (specifically, **Tables S6-S8** and **Figure S2)**.

### 3.1. Radiation characteristics

The 95 retrieved ROAs from Hawaii, Canary Islands, Galapagos and Fiji covered a large taxonomic range, including 30 orders and 64 families (**Figure 1A** and **Table S3**). They were largely dominated by invertebrates (60%: 58 ROAs) and plants (37%: 35 ROAs, all being vascular plants), with only three vertebrate ROAs, comprising two bird (Hawaiian Honeycreepers and Galápagos finches) and one reptile dataset (Galápagos Chelonoidis tortoises) **(Figure 1A)**. 73 ROAs (about 77%) had fewer than 50 described species. Conversely, only six had a hundred or more species, five being in Hawaii and one in the Canary Islands (**Figure 1B** and **Table S3**). Overall, the difference in ROA size between archipelagos was marginally non-significant (**Figure 1B**). The majority of ROA constituent species were SIE (median = 71%) and this proportion did not vary significantly between archipelagos (**Figure 1C**). For both Hawaii and the Canary Islands, where comparisons between plants and invertebrates were possible (**Figure 1D, E**), we found that invertebrate ROAs in Hawaii were on average larger and with a higher proportion of SIEs than plant ROAs, while no specific differences were found between taxa for the Canary Islands.

### 3.2. Univariate relationships between geo-environmental factors and Island S_ROA_

The results of the univariate GLMs before applying the FDR correction revealed significant linear relationships between Island S_ROA_ and elevation for about 44% of the ROAs, followed by inter-island isolation and area (both 39%), SDS (14%) and age (12%). We also identified 14% of ROAs showing a hump-shaped relationship with age fitting our criteria (**Figure 2**). Notably, most of the observed significant relationships with area, elevation, SDS and inter-island isolation had a sign aligned with our predictions (see a selection of relationships in **Figure 3**; all univariate plots are shown in **Supplementary Online Data**).

**Figure 2.**
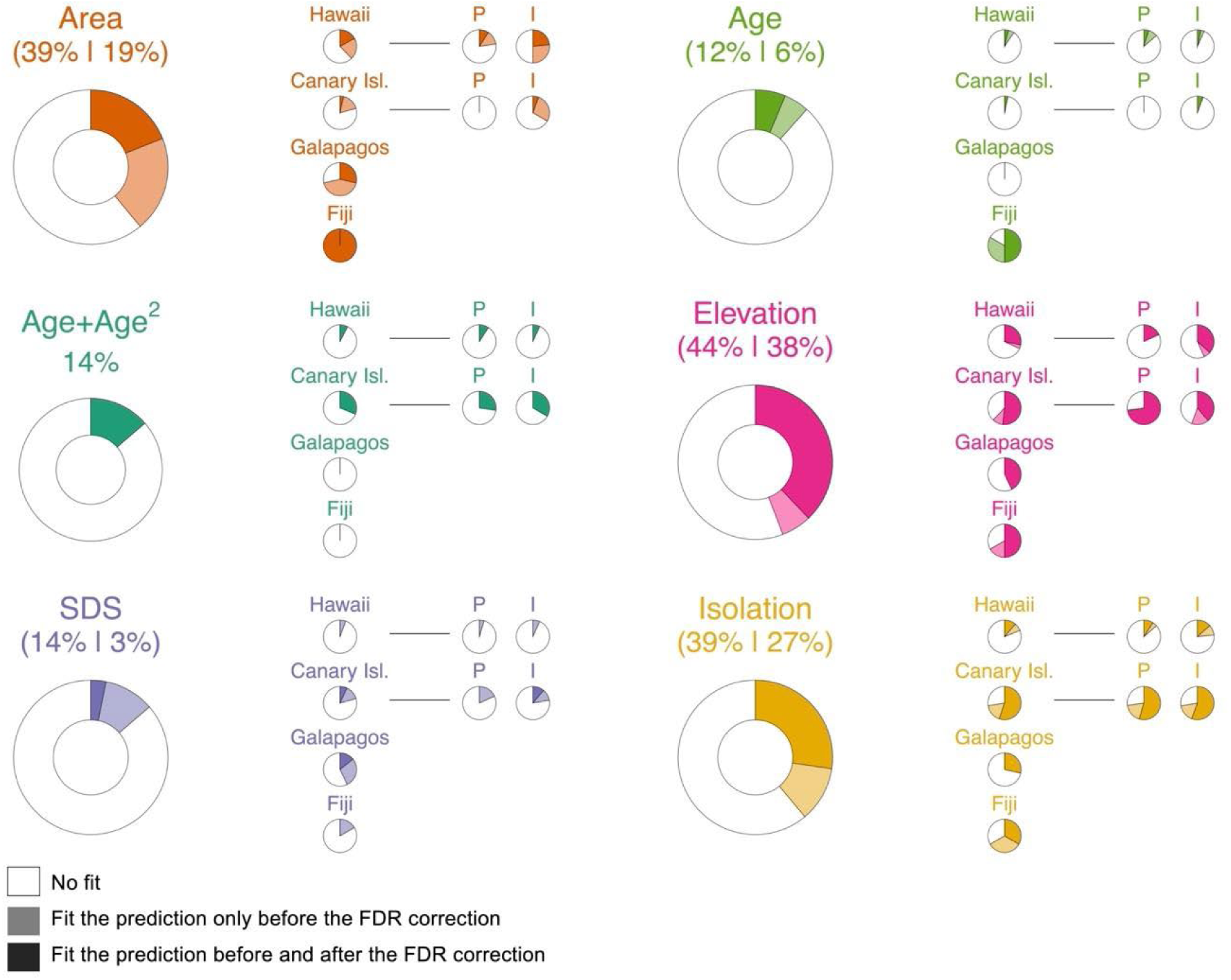
Summary of the results of the univariate GLMs between Island S_ROA_ and each geo-environmental factor separately. The coloured part within each pie chart shows the proportion of radiations having a significant relationship between Island S_ROA_ and a given factor. Because of multiple testing, the *P*-values of each individual GLM were corrected using the False Discovery Rate method (FDR, Benjamini & Hochberg 1995) to guard against inflation of Type-I errors. Therefore, the coloured part represents the proportion of relationships that were significant before FDR correction (the first number in parentheses above the pie chart) while the dark-shaded part shows the proportion of significant relationship that remained significant after FDR (false discovery rate) correction (i.e. second number in parentheses above the pie chart). The light-shaded part therefore represents the proportion of relationships that were significant only before FDR correction. Specifically for age + age^2^, only the proportion of radiations having a hump-shaped relationship with age that met our three criteria is shown (See Materials and Methods). For each factor, the four pie charts on the right show the proportion of significant relationships for each archipelago and for the plant and invertebrate subsets for Hawaii and the Canary Islands.

**Figure 3.**
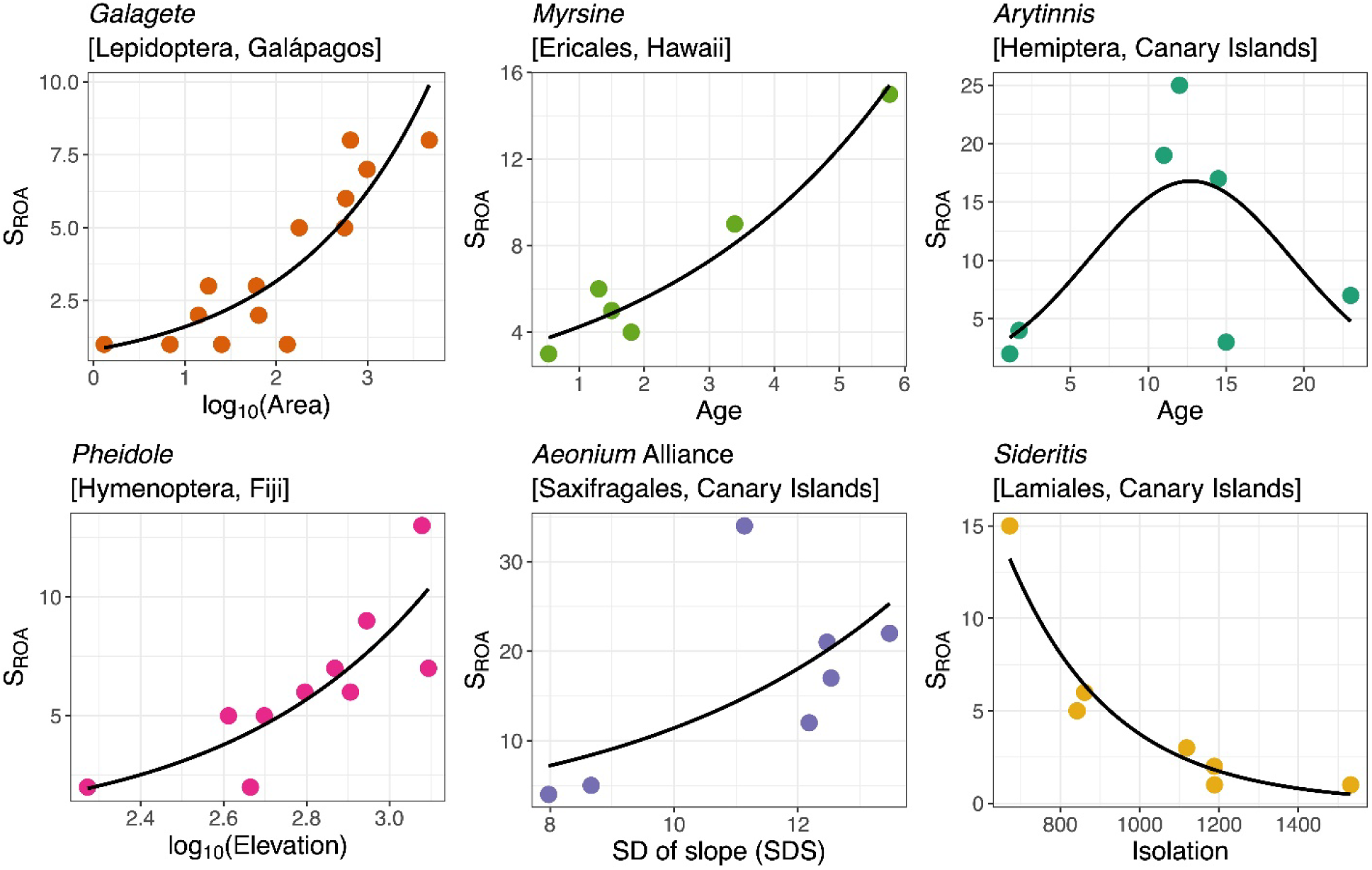
Selection of significant relationships between Island S_ROA_ (species richness of radiation on oceanic archipelago) and the geo-environmental factors. One radiation was selected for each factor, namely log_10_(area), age, age+age^2^, log_10_(elevation), SDS (standard deviation of slope) and isolation. Dots indicate islands and the black curves show the fitted values extracted from the Poisson / Negative binomial generalized linear model. All relationships were significant at 5% after the FDR correction. Plots for all ROAs and all factors are available in **Supplementary Online Data**.

Specifically, 100% of the significant relationships with area, 98% with elevation, and 70% with SDS were positive, while 92% of the significant relationships with inter-island isolation were negative. For age, eight of the 11 significant relationships recovered were positive. As importantly, the proportion of significant relationships between Island S_ROA_ and the factors appears to vary considerably from one archipelago to another (**Figure 2**). For example, in Hawaii, a higher proportion of ROAs exhibited significant relationships with area compared to the Canary Islands (37% and 20% of the relationships respectively), while all Fiji’s ROAs scaled significantly with area. Furthermore, the proportion of ROAs in the Canary Islands showing significant relationships with elevation was higher than in Hawaii (62% and 32% respectively). In addition, significant relationships with inter-island isolation and a hump-shaped relationship with age were more prevalent among ROAs in the Canary Islands than in all other archipelagos (**Figure 2**). In contrast, differences between taxa within each archipelago were less pronounced (**Figure 2**). We noticed a higher proportion of significant relationship with area for invertebrates in Hawaii and higher proportion of significant relationships with elevation for plants in the Canary Islands (**Figure 2**). Overall, similar trends were obtained after FDR correction of the *P*-values, but with a substantial decrease in the proportion of significant relationships from a 15% decrease for Elevation to 85% for SDS (**Figure 2**).

### 3.3. Testing whether the magnitude of the effect of geo-environmental factors on Island S_ROA_ varies between archipelagos and taxa

The differences observed between archipelagos in the proportion of significant relationships between factors and Island S_ROA_ were confirmed by the ANOVAs carried out to identify predictors of the strength of the effect of the factors on Island S_ROA_ (here quantified by the standardized β coefficient). Indeed, the ANOVAs explicitly demonstrated the importance of archipelago in explaining the strength of the effect of all factors on Island S_ROA_ (**see Table 1**), with post hoc tests revealing consistent pairwise differences between archipelagos as previously mentioned (see **Table S9**). To a lesser extent, the taxon also predicted the strength of the effects of area, SDS and isolation, with the effect of area and inter-island isolation being more pronounced for invertebrates than for plants while the opposite was true for SDS. Interestingly, we also found a negative effect of radiation size on the strength of effects of area and elevation, while the proportion of SIE exhibited a positive relationship with the same two factors (see **Table 1**). Similar outcomes were obtained when we conducted the analyses with a subset including only Hawaii and the Canary Islands and restricted the analyses to plants and invertebrates (**Table S9**). As importantly, these last analyses unveiled significant interactions between archipelagos and taxa for age, elevation and isolation, supporting the fact that the difference between taxa in the effect of these factors was archipelago dependent (see **Table 1** and **Table S9**).For instance, for Canarian ROAs, the effect of elevation was stronger for plants than for invertebrates but did not differ between the two taxa in Hawaii (**Table S9**).

**Table 1.** Results of the ANOVAs testing whether the strength of the effect of the geo-environmental factors on the Island S_ROA_ can be explained by archipelago, taxa, radiation size (log_10_-tranformed), and proportion of SIE. The effect of a given geo-environmental factor on the Island S_ROA_ of a given ROA is quantified by its standardized β coefficient for the factors area, age, elevation, standard deviation of slope (SDS) and inter-island isolation while the effect of age + age^2^ is coded as a binary variable with a value of 1 if S_ROA_ follows a hump-shaped relationship with age according to our three criteria (See main text) and 0 if not. For the age + age^2^ model, the model is fitted with a binomial GLM. The R^2^ is given for each model and the *P*-value is provided for each predictor remaining in the model after the backward elimination procedure. Analyses were initially performed for all radiations without testing the interaction between Archipelago and Taxa owing to the design being unbalanced. The models with the interaction term were performed for a subset of the data including only the radiations of Hawaii and the Canary Islands and for plants and invertebrates. Note that archipelago and taxa were kept in the model when their interaction was significant even if their independent effect was not. Results of the post-hoc tests are given in **Table S9**. For the predictors Size and Proportion of SIE, the sign of the relationship is given in parentheses. Significant effects are marked in bold.

| | | Predictors | | | | | $R^2$ |
| --- | --- | --- | --- | --- | --- | --- | --- |
|  | Factors | Archipelago | Taxa | A x T | Radiation Size | Proportion of SIE |  |
| All radiations |  |  |  |  |  |  |  |
| Standardized $\beta$ coefficient | Area | **<0.001** | **0.026** | - | **<0.001 (-)** | **<0.001 (+)** | 0.541 |
|  | Age | **0.031** |  | - |  |  | 0.090 |
|  | Age + age<sup>2</sup> | **0.001** |  | - |  |  | 0.288 |
|  | Elevation | **<0.001** |  | - | **<0.001 (-)** | **0.007 (+)** | 0.425 |
|  | SDS | **<0.001** | **<0.001** | - |  |  | 0.527 |
|  | Inter-island isolation | **<0.001** | **0.018** | - |  |  | 0.227 |
| Subset |  |  |  |  |  |  |  |
| Standardized $\beta$ coefficient | Area | | **0.037** | | **<0.001 (-)** | **0.004 (+)** | 0.231 |
|  | Age | 0.561 | 0.901 | **0.003** |  | **0.031 (+)** | 0.138 |
|  | Age + age<sup>2</sup> | **0.007** |  |  |  |  | 0.146 |
|  | Elevation | **0.015** | 0.730 | **<0.001** | **<0.001 (-)** |  | 0.318 |
|  | SDS | **0.003** | **<0.001** |  |  |  | 0.267 |
|  | Inter-island isolation | **<0.001** | **0.044** | **0.034** |  |  | 0.314 |

### 3.4 Quantifying the relative contribution of the geo-environmental factors in explaining Island S_ROA_ variations between archipelagos and taxa

Figure 4 presents the results of the GLMMs (run separately for each archipelago) including factor contribution (Figure 4A), the R^2^ (Figure 4B) and averaging standardized estimates (Figure 4C). The GLMMs shed further light on the role of the archipelago and the taxon in relation to the effects of geo-environmental factors on Island S_ROA_. For Hawaii, we recovered a positive effect of area and SDS, with an averaged R^2^m (fixed effects only) accounting for approximately 40% of the variation in Island S_ROA_. For the Canary Islands, best models explained 32% of the variance (R^2^m) and included a humped-shaped relationship with age, a positive effect of elevation and a negative effect of isolation. In contrast, no specific factor emerged as significant in the Galápagos GLMMs and the model selection procedure exhibited high uncertainty (**Table S10**) with only a small fraction of the variance explained (R^2^m = 19%). For Fiji, we identified a humped-shape relationship with age and a positive effect of elevation with an R^2^m of 70%. When performed solely for plant ROAs, the analyses highlighted a strong positive effect of elevation for both Hawaii and the Canary Islands while a humped-shape relationship with age was detected for Hawaiian plant ROAs. Overall, the plant ROA models exhibited a R^2^m of 46% for both archipelagos. For invertebrate ROAs, the results for Hawaii emphasised the importance of area and SDS with an average R^2^m of 50%, while for the Canary Islands, humped-shape relationship with age, a positive effect of elevation and a negative effect of inter-island isolation were identified with an average R^2^m of 30%. Importantly, for most archipelagos and subsets, the variance captured by the random structure was high, reaching 47% for Galápagos. Further details of these analyses, including the best random structures and family distribution are given in **Table S6.**

**Figure 4.**
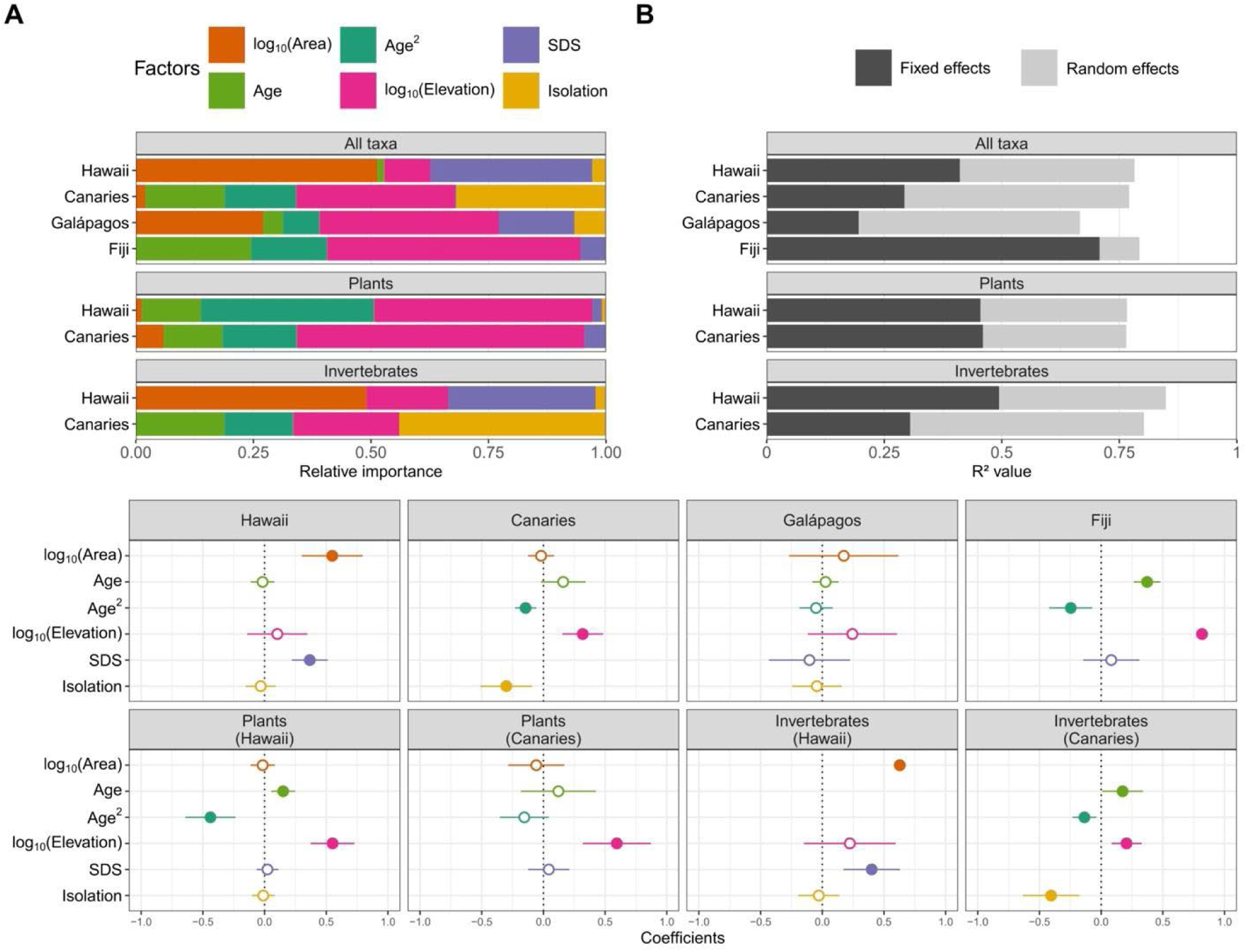
(A) The relative importance of each fixed effect as a predictor of Island S_ROA_ for all ROAs in each archipelago, followed by the plant and invertebrate subsets for Hawaii and the Canary Islands; (B) decomposition of the variance explained by the best set of models (conditional R^2^c) into its components assigned to the fixed effects (marginal R^2^m, dark grey) and random effects (R^2^c - R^2^m, light grey) for each subset; (C) standardized coefficients of the effect of the factors on Island S_ROA_ variation and their associated confidence intervals, extracted from the model constructed by averaging over the best models identified for each archipelago and for the plant and invertebrate subsets. Solid dots indicate a significant effect, while empty dots indicate no effect. Details for the best models are in Table S10.

## 4. Discussion

Despite the large amount of work undertaken on ROAs in recent decades, little research has focused on the emergent diversity patterns, with the few studies that have done so focusing on the effect of island area and geological age (e.g. Lim & Marshall 2017, Gillespie & Baldwin 2010; but see Parent & Crespi 2006 and Roell et al. 2021). In this study, we have examined how the Island S_ROA_ of almost a hundred ROAs scale with five major geo-environmental factors (area, age, elevation, topographic complexity and intra-archipelago isolation), and whether the magnitude of their effects was archipelago and taxon specific.

### 4.1 A brief overview of the selected ROAs

Our dataset reveals a clear dominance of ROAs from Hawaii and the Canary Islands in the literature that qualify for our basic criteria for inclusion (e.g. purely volcanic, oceanic islands; minimum of 6 islands, etc.). This was unsurprising since these two archipelagos are known to host a tremendous number of successful ROAs (e.g., Machado et al. 2017, Hembry et al. 2021, Cerca et al. 2023). These archipelagos are also two of the best studied volcanic island systems in the world, both in terms of their natural history and the phylogenetic origin of their biota (e.g., Caujapé-Castells et al. 2017, Price & Wagner 2018). On the other hand, the Galápagos, which are more ecologically uniform and geologically younger (Naciri & Linder 2020), are home to smaller ROAs, many of which number fewer than 10 species (Parent et al. 2008) and were thus excluded from our study. The few ROAs retrieved for Fiji can be explained by the overall less advanced phylogenetic understanding of Fijian endemic groups compared to the other three archipelagos (Rudbeck et al. 2022).

The study of evolutionary radiations invariably raises the question of whether they are adaptive or not. Within island ROAs it appears that non-adaptive speciation mechanisms may hold considerable relevance, notwithstanding the evidence of trait shifts and trait disparification within particular ROAs (see, e.g. Dimitrov et al. 2008, Benavides et al. 2008, Groom et al. 2013, Vitales et al. 2015, Chiba & Cowie 2016, Schenk 2021, Barajas-Barbosa et al. 2023, Cerca et al. 2023, Whittaker et al. 2023, Illera et al. 2024). However, robust attribution of the mode of speciation, the relevance of adaptive and non-adaptive mechanisms, and of allopatric vs sympatric episodes tends to be lacking is generally lacking for the systems considered herein and therefore we therefore made no attempt to sub-divide the ROAs by such categorisations in the present study (See Hernández-Hernández 2019).

### 4.2. Island geo-environmental factors, and specifically elevation, are effective drivers of Island S_ROA_

Our study confirms our first hypothesis that the selected geo-environmental factors were overall good predictors of Island S_ROA_, but they were not all equally important. Indeed, most of the significant relationships recovered were attributed, in a decreasing order, to elevation, inter-island isolation and area with a sign in line with our predictions, i.e. positive for area and elevation and negative for inter-island isolation. In this respect, these results corroborate what is usually reported in studies on island species richness of larger groups (vascular plants, Kreft et al. 2008; spiders, Cardoso et al. 2010; birds, Kalmar & Currie 2006). The predominance of elevation in our analyses confirms our prediction that environmental heterogeneity is a key driver of Island S_ROA_. This is further supported by the high proportion of SIE detected in the ROAs (median 71%), confirming the role of between-island (i.e. where different SIE occur on different islands in an archipelago) and within-island speciation (where multiple SIE occur within a single island) and underscoring the importance of environmental heterogeneity in the radiation process (Cowie & Holland 2008). The elevation gradient often serves as a proxy of habitat diversity and is closely associated with adaptive processes in ROA diversification (e.g., Givnish 1997, Schluter 2000). Indeed, several radiated species in many different ROAs have developed into locally adapted ecotypes in response to elevational gradients (Jorgensen & Olesen 2001 and see example below). High-elevation zones can also provide refugia during periods of climate change, allowing relict populations of ancient species to persist (Irl et al. 2015). Elevation can also act as a direct surrogate of topographic complexity, as high elevation is a prerequisite for the presence of a complex landscape structure promoting within-island allopatry (Brown et al. 2006). In this context, it is worth highlighting that the ROAs exhibiting a strong relationship between Island S_ROA_ and elevation were also typically the ones with the highest proportion of SIE (**Table 1**). Note that the SDS factor, which quantifies topographical complexity *per se* in our analyses, exhibited very few significant relationships with Island S_ROA_. This was somewhat anticipated, since its effect seems to become apparent only after controlling for other factors such as area or age, as reported by Roell et al. (2021) and Barajas-Barbosa et al. (2020). Most significant relationships between Island S_ROA_ and inter-island isolation revealed that, as expected, the most remote islands in the archipelago were the least diverse. However, inter-island isolation did not affect Island S_ROA_ for 60% of the ROAs. In some cases, the effects of inter-island isolation may have been obscured by subsequent within-island allopatry or back-colonization following inter-island speciation (Cowie & Holland 2008). Additionally, the high vagility of certain ROA species may have prevented geographic speciation between islands, with radiation in these cases resulting primarily from sympatric speciation (Losos & Ricklefs 2009). It is important to note that we focused on a single scale of isolation—island isolation within the archipelago—because our primary goal was to test the effect of island-specific characteristics on Island S_ROA_. However, we should not exclude the possibility that isolation may operate at other scales. Isolation of the archipelago itself could indirectly impact Island S_ROA_ by affecting the overall size of the ROA (Tao et al. 2021). Within-island isolation, i.e., the isolation between populations on the same island, will also play a crucial role in determining Island S_ROA_ by driving allopatric speciation, although we expect topographic complexity to capture such mechanisms to some extent (Barajas-Barbosa et al. 2020).

The weak effect of age on Island S_ROA_, whether considered through a linear or hump-shaped relationship, may stem from the fact that age tends to have its strongest influence when interacting with area (Whittaker et al. 2008). This aligns with the idea that the impact of time on speciation cannot be fully decoupled from the amount of ecological space available for it as postulated by the GDM (Whittaker et al. 2008, Borregaard et al. 2017). Moreover, our results may also show the limitation of assigning a single age to an island, given that many of the oceanic islands we considered are formed by different basaltic shields of varying ages (Irl et al. 2015, Otto et al. 2016). It may also be noted that for much of the last 1M years, the islands of Lanzarote and Fuerteventura have been united by lower eustatic sea-levels to form a single island, thus arguably conveying the biological legacies of Fuerteventura’s greater age (23 Ma) on the island of Lanzarote (15 Ma). Were the age of 23 Ma used for both islands this would make a stronger case for a humped-relationship between Island SROA. It is also important to note that our measure of island age reflects the age of the platform within which ROAs have developed, but takes no account of the actual colonization dates of the founding lineages from which each ROA has developed. As we gain tighter estimates of the colonization date of more ROAs it may become possible to take time since arrival directly into account in future modelling exercises.

Finally, the above results should be viewed with caution due to the substantial correlation between geo-environmental variables observed in most of the archipelagos studied (Figure S1), which may lead to confounding effects.

### 4.3. Archipelago and taxon determine the strength of the effect of geo-environmental factors on Island S_ROA_

The Island S_ROA_ of a ROA on a particular island is the outcome of many processes affecting the colonization, evolution, and persistence of island populations. One important finding of our study is to show that the expression of these processes is likely to differ among archipelagos because of differences in their geo-environmental dynamics (e.g., island chronosequence, habitat diversity, spatial configuration) but also among taxa due to the variation in biological attributes among ROAs (e.g., dispersion, adaptive capacity, population density, time of colonization) (**Table 1** and Figure 2 and **4**). While other authors have already highlighted an archipelagic and/or taxonomic dependence in the factors driving diversity patterns of ROAs (e.g., Parent 2012, Lim & Marshall 2017), our study is the first to quantify such effects across multiple archipelagos and taxa via a comparative statistical approach.

Taken together, variation in Island S_ROA_ of Hawaiian ROA were mostly driven by area and topographic complexity (SDS). The explanation for this likely lies in the role of ‘the Big Island’ in driving diversity patterns in Hawaii. Indeed, our results and several other studies (e.g. Price & Wagner 2011, Eckstut et al. 2011) show that for most ROAs, the Big Island has a diversity value lower than predicted by the ISAR with the other Hawaiian Islands (**Supplementary Online Data**), reflecting that the island is both the largest and the youngest island (0.53 Ma, **Table S4**) in origin. In our case, this deviation is captured by the comparatively lower topographic complexity of the Big Island, indicative of its smooth and young terrain, compared to the older and more rugged islands (Price & Wagner 2011, Barajas-Barbosa et al. 2020). Given that Hawaii hosts some of the largest ROAs, and with the highest proportion of SIEs (Figure S3), we cannot rule out that the positive effect of SDS on Island S_ROA_ may reflect the importance of species accumulation through within-island allopatry during the radiating process (Carson & Kaneshiro 1976, Hembry et al. 2021). In contrast to Hawaii, isolation, elevation and island age (hump-shaped) appear to shape the Island S_ROA_ patterns for the Canary Islands (Figure 4). The predominance of a negative effect of inter-island isolation reveals that species accumulation was higher in the central, least isolated islands within the archipelago, namely Tenerife and Gran Canaria. This aligns with previous studies that have positioned these two islands as primary centers of diversification for the extant species of the archipelago (e.g. Francisco-Ortega et al. 2002, Allan 2004, Fernández-Mazuecos & Vargas 2011), making them the main sources of dispersal for the peripheral western islands. Using a Bayesian framework to infer dispersion processes within the archipelago, Sanmartín et al. (2008) indeed identified a stepwise dispersal pattern from the central islands to the islands of La Palma and El Hierro in the west. This distinctive feature of these two central islands relies also on their intrinsic characteristics, such as their elevation (3,715 and 1,956 m.a.s.l respectively) and topographic complexity, both resulting from their intermediate age (12 and 14.5 Myr respectively) relative to the age of the archipelago (i.e., 23 Myr). According to the GDM, intermediate-age islands reach ’maturity’ through the slowing down of island-building processes in favour of erosive processes, generating a more complex topography (Borregaard et al. 2017). In that respect, we found that two-thirds of the ROAs of Canary Islands had their highest values of Island S_ROA_ and SIE in Tenerife (see **Figure S4**). This relationship with age was reflected in the importance of the hump-shaped relationship detected for the Canary Islands (Figure 4), with young and old islands having low Island S_ROA_ values compared to the central ones. However, the low Island S_ROA_ of the two oldest islands (i.e., Lanzarote and Fuerteventura) are not simply or exclusively the result of the island aging process as predicted by the GDM. It also reflects that these islands underwent climatic fluctuations during the Pleistocene followed by a recent onset of an arid climate that wiped out species not adapted to these novel conditions, thereby reinforcing the hump-shaped relationship with age (García-Verdugo et a. 2019).

For both Hawaii and the Canary Islands, the two archipelagos for which a comparison between vascular plants and invertebrates ROAs was possible, taxon also appears to wield a significant role in driving the effect of the geo-environmental factors on Island S_ROA_. Probably the most striking finding here was the strong effect of elevation detected for the plant subset of both archipelagos. Indeed, several emblematic plant adaptive ROAs span the entire elevation gradient of islands, with different species each adapted to narrow habitats, ranging from dry coastal areas to mesic environments and wet forests [e.g. Lobeliads (Givnish et al. 2009), Silversword alliance (Baldwin & Sanderson 1998), *Schiedea* (Sakai et al. 2006) and *Euphorbia* (Yang et al. 2018) in Hawaii; *Aeonium* Alliance (Kim et al. 2008, Messerschmid et al. 2023), *Argyranthemum* (White et al. 2020), *Echium* (García-Maroto et al. 2009), *Sonchus* (Kim et al. 2008) in the Canary Islands]. In contrast, the relative contribution of elevation for invertebrates (Figure 4) was very limited in both archipelagos. Several studies suggest that diversification in many arthropod ROAs on oceanic islands has been primarily driven by niche partitioning resulting from biotic interactions, such as microhabitat choice or host plant specialization, rather than by the island ecological/climatic gradients (Bennet & O’Grady 2012, Machado et al. 2017, Johns et al. 2018, Cotoras et al. 2018). A recent study on Hawaiian radiations even indicated that strong environmental conservatism dominates for arthropod ROAs as they largely occur within restricted environmental envelopes (Hiller et al. 2019, Dorey et al. 2020).

The best models for Fiji highlighted the pivotal roles of age and elevation in shaping the diversification of Fijian ROAs. Firstly, all Fijian ROAs included in our study reached their maximum Island S_ROA_ on Viti Levu, the oldest island of Fiji (**Figures S4** and **Supplementary Online Data**). The observed effect of age + age² detected in our GLMM was mainly due to a non-linear increase of Island S_ROA_ with age, rather than a true hump-shaped relationship (Figure 4 and **Supplementary Online Data**). Secondly, studies examining the presence of a taxon cycle in Fijian ants, in particular the *Strumigenys* and *Pheidole* ROAs, supported the hypothesis that the radiation process was primarily associated with niche shifts towards higher elevations (Sarnat & Economo 2012, Liu et al. 2020), with species accumulation being more pronounced on high-elevation islands. Moreover, diversification of *Cyrtandra* in Fiji is thought to have been triggered by the pronounced elevational gradient with several small-range species of the rainforest understory being distributed from the lowlands to the montane zone (Johnson 2017, 2023). Conversely, the absence of significant signal for Galápagos ROAs could be due to the wide taxonomic range covered by the seven retrieved ROAs (two beetles, one moth, one bird, one reptile, one land snail and one flowering plant ROA), which might have hindered the identification of common drivers.

Finally, it should be noted that the random structure of our GLMMs, mostly the random effect of radiation identity for slope and intercept (**Table S6**), occasionally demonstrated an explanatory power greater than the geo-environmental factors in explaining Island S_ROA_ variation (Figure 4B and **Table S6**). Analytically, this suggests that while a given geo-environmental factor may influence most radiations similarly within an archipelago (e.g. a positive relationship between elevation and Island S_ROA_ in the Canary Islands, Figure 4), the magnitude of its effect can vary significantly between radiations. This finding is important because this variance likely reflects finer differences between ROAs, particularly in terms of their ecological and evolutionary characteristics (e.g., dispersal abilities, speciation rates), which are not accounted for by our fixed effects. Overall, this highlights the need for a deeper understanding of how ROA life history traits and geo-environmental factors interact to influence the radiation process.

### 4.4. Limitations

Our approach undoubtedly comes with some limitations. First, as detailed above, our study is clearly biased toward Hawaii and the Canary Islands and, as such, may not fully represent the diversity of oceanic archipelagos and ROAs worldwide. Second, our values of Island S_ROA_ may be potentially biased due to unrecorded human mediated species extinctions or local (islands) extirpations (Paulay 1994, Lim & Marshall 2017, Whittaker et al. 2023). Moreover, we cannot exclude the possibility that inconsistencies among taxonomists and the varying degrees of completeness of these endemic clades may have generated noise in our analyses. However, the analyses that included subspecies did not show any specific differences with the analyses run without subspecies, suggesting that our results may be robust to marginal variation in species number (**Table S7, S8** and **Figure S2**). Third, we acknowledge that the limited number of islands within each archipelago poses a challenge in terms of statistical power and restricts the use of complex models. Fourth, the ten species threshold we chose to ensure sufficient variance in Island S_ROA_ between islands introduces a bias toward “successful” ROAs, and our conclusions therefore might not apply for more modest ROAs.

### 4.5. Conclusion & Perspectives

Our study confirms that geo-environmental factors are effective predictors of Island S_ROA_, aligning with previous biogeographic expectations. Therefore, it seems that the main abiotic forces driving overall island biodiversity also govern the diversification of endemic clades. Elevation emerged as a key factor, complementing existing theoretical and empirical research that has highlighted the central role of environmental heterogeneity in driving diversification on oceanic islands (e.g., Cabral et al. 2019, Roell et al. 2021). The influence of geo-environmental factors also varied by archipelago and taxon, indicating that unique archipelagic dynamics and biological characteristics channel diversification differently. Future analyses should explore the contribution of additional factors such as climate change and variability (e.g., Barajas-Barbosa et al. 2020) as well as direct measurements of habitat diversity (Yu et al. 2015). Moreover, accounting for palaeo-configuration (Norder et al. 2018) may be particularly relevant in the context of ROAs, as variation in archipelago configuration over geological time is likely to have left detectable evolutionary legacies in contemporary distributions of endemism (Weigelt et al. 2016, Fernández-Palacios et al. 2016).

## Acknowledgments

We thank Delphine Montagne for her help with QGIS. We also thank the organisers of the Island Biology 2023 conference in Lipari, which enabled us to have fruitful live discussions on this work, and we thank the journal reviewers for their constructive feedback on our initial submission.

## Author Contributions

B.B. and F.R. designed the study. K.A.T. helped compile the list of oceanic archipelagos. B.B. collected the data on evolutionary radiations and F.R. collected the data on island characteristics. B.B. and F.R. analysed the data. B.B. and F.R. drafted the manuscript in collaboration with T.J.M., J.M.F.P, C.P., K.A.T. and R. J.W.. All authors contributed to and approved the submitted versions.

## Data and code availability

All the data and code used in this manuscript are available in Supplementary Information and in the Supplementary Online Data available in the following repository: 10.5281/zenodo.15680633.

## References

1. Allan, G. J., Francisco-Ortega, J., Santos-Guerra, A., Boerner, E., & Zimmer, E. A. (2004). Molecular phylogenetic evidence for the geographic origin and classification of Canary Island *Lotus* (Fabaceae: Loteae). Molecular Phylogenetics and Evolution, 32(1), 123–138.

2. Amorim, I. R., Emerson, B. C., Borges, P. A. V., & Wayne, R. K. (2012). Phylogeography and molecular phylogeny of Macaronesian island *Tarphius* (Coleoptera: Zopheridae): Why are there so few species in the Azores? Journal of Biogeography, 39(9), 1583–1595. 10.1111/j.1365-2699.2012.02721.x

3. Baldwin, B. G., & Sanderson, M. J. (1998). Age and rate of diversification of the Hawaiian silversword alliance (Compositae). Proceedings of the National Academy of Sciences, USA, 95(16), 9402–9406.

4. Barajas Barbosa, M. P., Craven, D., Weigelt, P., Denelle, P., Otto, R., Díaz, S., … Kreft, H. (2023). Assembly of functional diversity in an oceanic island flora. Nature, 619(7970), 545–550.

5. Barajas-Barbosa, M. P., Weigelt, P., Borregaard, M. K., Keppel, G., & Kreft, H. (2020). Environmental heterogeneity dynamics drive plant diversity on oceanic islands. Journal of Biogeography, 47(10), 2248–2260. 10.1111/jbi.13925

6. Bartoń, K. (2023). MuMIn : Multi-Model Inference. R package version 1.47.5, <https://CRAN.R-project.org/package=MuMIn>.

7. Beatty, C. D., Sánchez Herrera, M., Skevington, J. H., Rashed, A., Van Gossum, H., Kelso, S., & Sherratt, T. N. (2017). Biogeography and systematics of endemic island damselflies: The *Nesobasis* and *Melanesobasis* (Odonata: Zygoptera) of Fiji. Ecology and Evolution, 7(17), 7117–7129. 10.1002/ece3.3175

8. Benavides, E., Baum, R., Snell, H. M., Snell, H. L., & Sites, Jr., J. W. (2008). Island biogeography of Galápagos lava lizards (Tropiduridae : *Microlophus*): species diversity and colonization of the archipelago. Evolution, 63(6), 1606–1626. 10.1111/j.1558-5646.2009.00617.x

9. Benjamini, Y., & Hochberg, Y. (1995). Controlling the false discovery rate: A practical and powerful approach to multiple testing. Journal of the Royal statistical society: series B (Methodological), 57(1), 289–300.

10. Bennett, G. M., & O’Grady, P. M. (2012). Host–plants shape insect diversity: Phylogeny, origin, and species diversity of native Hawaiian leafhoppers (Cicadellidae: *Nesophrosyne*). Molecular Phylogenetics and Evolution, 65(2), 705–717. 10.1016/j.ympev.2012.07.024

11. Borges, P. A. V., & Hortal, J. (2009). Time, area and isolation: Factors driving the diversification of Azorean arthropods. Journal of Biogeography, 36(1), 178–191. 10.1111/j.1365-2699.2008.01980.x

12. Borregaard, M. K., Amorim, I. R., Borges, P. A. V., Cabral, J. S., Fernández-Palacios, J. M., Field, R., … Whittaker, R. J. (2017). Oceanic island biogeography through the lens of the general dynamic model: Assessment and prospect. Biological Reviews, 92(2), 830–853. 10.1111/brv.12256

13. Brooks, M. E., Kristensen, K., Van Benthem, K. J., Magnusson, A., Berg, C. W., Nielsen, A., … Bolker, B. M. (2017). glmmTMB balances speed and flexibility among packages for zero-inflated generalized linear mixed modeling. The R journal, 9(2), 378–400.

14. Brown, R. M., Siler, C. D., Oliveros, C. H., Esselstyn, J. A., Diesmos, A. C., Hosner, P. A., … Alcala, A. C. (2013). Evolutionary processes of diversification in a model island archipelago. Annual Review of Ecology, Evolution, and Systematics, 44(1), 411–435. 10.1146/annurev-ecolsys-110411-160323

15. Brown, R. P., Hoskisson, P. A., Welton, J.-H., & Baez, M. (2006). Geological history and within-island diversity: A debris avalanche and the Tenerife lizard *Gallotia galloti*. Molecular Ecology, 15(12), 3631–3640.

16. Cabral, J. S., Whittaker, R. J., Wiegand, K., & Kreft, H. (2019). Assessing predicted isolation effects from the general dynamic model of island biogeography with an eco-evolutionary model for plants. Journal of Biogeography, jbi.13603. 10.1111/jbi.13603

17. Cardoso, P., Arnedo, M. A., Triantis, K. A., & Borges, P. A. V. (2010). Drivers of diversity in Macaronesian spiders and the role of species extinctions: Diversity of Macaronesian spiders. Journal of Biogeography, 37(6), 1034–1046. 10.1111/j.1365-2699.2009.02264.x

18. Carson, H. L., & Kenneth, K. Y. (1976). *Drosophila* of Hawaii: Systematics and Ecological Genetics. Annual Review of Ecology and Systematics, 311–345. 10.1146/annurev.es.07.110176.001523

19. Caujapé-Castells, J., García-Verdugo, C., Marrero-Rodríguez, Á., Fernández-Palacios, J. M., Crawford, D. J., & Mort, M. E. (2017). Island ontogenies, syngameons, and the origins and evolution of genetic diversity in the Canarian endemic flora. Perspectives in Plant Ecology, Evolution and Systematics, 27, 9–22. 10.1016/j.ppees.2017.03.003

20. Cerca, J., Cotoras, D. D., Bieker, V. C., De-Kayne, R., Vargas, P., Fernández-Mazuecos, M., … Martin, M. D. (2023). Evolutionary genomics of oceanic island radiations. Trends in Ecology & Evolution, 38(7), 631–342. 10.1016/j.tree.2023.02.003

21. Chiba, S., & Cowie, R. H. (2016). Evolution and extinction of land snails on oceanic islands. Annual Review of Ecology, Evolution, and Systematics, 47(1), 123–141. 10.1146/annurev-ecolsys-112414-054331

22. Cotoras, D. D., Bi, K., Brewer, M. S., Lindberg, D. R., Prost, S., & Gillespie, R. G. (2018). Co-occurrence of ecologically similar species of Hawaiian spiders reveals critical early phase of adaptive radiation. BMC Evolutionary Biology, 18(1), 100. 10.1186/s12862-018-1209-y

23. Cowie, R. H., & Holland, B. S. (2008). Molecular biogeography and diversification of the endemic terrestrial fauna of the Hawaiian Islands. Philosophical Transactions of the Royal Society B: Biological Sciences, 363(1508), 3363–3376. 10.1098/rstb.2008.0061

24. Darwin, C. (1845). Journal of researches into the natural history and geology of the countries visited during the voyage of HMS Beagle round the world (Vol. 7). London: John Murray.

25. de la Harpe, M., Paris, M., Karger, D. N., Rolland, J., Kessler, M., Salamin, N., & Lexer, C. (2017). Molecular ecology studies of species radiations: Current research gaps, opportunities and challenges. Molecular Ecology, 26(10), 2608–2622. 10.1111/mec.14110

26. Dimitrov, D., Arnedo, M. A., & Ribera, C. (2008). Colonization and diversification of the spider genus *Pholcus* Walckenaer, 1805 (Araneae, Pholcidae) in the Macaronesian archipelagos: Evidence for long-term occupancy yet rapid recent speciation. Molecular Phylogenetics and Evolution, 48(2), 596–614. 10.1016/j.ympev.2008.04.027

27. Dorey, J. B., Groom, S. V. C., Freedman, E. H., Matthews, C. S., Davies, O. K., Deans, E. J., … Schwarz, M. P. (2020). Radiation of tropical island bees and the role of phylogenetic niche conservatism as an important driver of biodiversity. Proceedings of the Royal Society B: Biological Sciences, 287(1925), 20200045. 10.1098/rspb.2020.0045

28. Eckstut, M. E., McMahan, C. D., Crother, B. I., Ancheta, J. M., McLennan, D. A., & Brooks, D. R. (2011). PACT in practice: Comparative historical biogeographic patterns and species-area relationships of the Greater Antillean and Hawaiian Island terrestrial biotas: Comparative PACT analyses. Global Ecology and Biogeography, 20(4), 545–557. 10.1111/j.1466-8238.2010.00626.x

29. Economo, E. P., Klimov, P., Sarnat, E. M., Guénard, B., Weiser, M. D., Lecroq, B., & Knowles, L. L. (2015). Global phylogenetic structure of the hyperdiverse ant genus *Pheidole* reveals the repeated evolution of macroecological patterns. Proceedings of the Royal Society B: Biological Sciences, 282(1798), 20141416. 10.1098/rspb.2014.1416

30. Emerson, B. C. (2002). Evolution on oceanic islands: Molecular phylogenetic approaches to understanding pattern and process. Molecular ecology, 11(6), 951–966. 10.1046/j.1365-294X.2002.01507.x

31. Fernández-Mazuecos, M., & Vargas, P. (2011). Genetically depauperate in the continent but rich in oceanic islands: *Cistus monspeliensis* (Cistaceae) in the Canary Islands. PLoS ONE, 6(2), e17172. 10.1371/journal.pone.0017172

32. Fernández-Palacios, J. M., Rijsdijk, K. F., Norder, S. J., Otto, R., Nascimento, L. de, Fernández-Lugo, S., … Whittaker, R. J. (2016). Towards a glacial-sensitive model of island biogeography. Global Ecology and Biogeography, 25(7), 817–830. 10.1111/geb.12320

33. Florencio, M., Patiño, J., Nogué, S., Traveset, A., Borges, P. A. V., Schaefer, H., … Santos, A. M. C. (2021). Macaronesia as a fruitful arena for ecology, evolution, and conservation biology. Frontiers in Ecology and Evolution, 9, 718169. 10.3389/fevo.2021.718169

34. Francisco-Ortega, J., Fuertes-Aguilar, J., Kim, S., Santos-Guerra, A., Crawford, D. J., & Jansen, R. K. (2002). Phylogeny of the Macaronesian endemic *Crambe* section *Dendrocrambe* (Brassicaceae) based on internal transcribed spacer sequences of nuclear ribosomal DNA. American Journal of Botany, 89(12), 1984–1990. 10.3732/ajb.89.12.1984

35. Funk, V. A., & Wagner, W. L. (1995). Biogeographic patterns in the Hawaiian Islands. In W. L. Wagner & V. A. Funk (Éds.), Hawaiian biogeography: Evolution on a hot spot archipelago. Smithsonian Institution Press.

36. García-Maroto, F., Mañas-Fernández, A., Garrido-Cárdenas, J. A., Alonso, D. L., Guil-Guerrero, J. L., Guzmán, B., & Vargas, P. (2009). Δ6-Desaturase sequence evidence for explosive Pliocene radiations within the adaptive radiation of Macaronesian *Echium* (Boraginaceae). Molecular Phylogenetics and Evolution, 52(3), 563–574. 10.1016/j.ympev.2009.04.009

37. García-Verdugo, C., Caujapé-Castells, J., Illera, J. C., Mairal, M., Patiño, J., Reyes-Betancort, A., & Scholz, S. (2019). Pleistocene extinctions as drivers of biogeographical patterns on the easternmost Canary Islands. Journal of Biogeography, 46(5), 845–859. 10.1111/jbi.13563

38. Gillespie, R. G. (2016). Island time and the interplay between ecology and evolution in species diversification. Evolutionary Applications, 9(1), 53–73. 10.1111/eva.12302

39. Gillespie, R. G., & Baldwin, B. G. (2010). Island biogeography of remote archipelagoes. In The theory of island biogeography revisited (p. 358–387). Princeton University Press Princeton.

40. Gillespie, R. G., Bennett, G. M., De Meester, L., Feder, J. L., Fleischer, R. C., Harmon, L. J., … Wogan, G. O. U. (2020). Comparing Adaptive Radiations Across Space, Time, and Taxa. Journal of Heredity, 111(1), 1–20. 10.1093/jhered/esz064

41. Givnish, T. J. (1997). Adaptive radiation and molecular systematics: Issues and approaches. In T. J. Givnish & K. J. Systema (Éds.), Molecular evolution and adaptive radiation (p. 1-54). Cambridge University Press.

42. Givnish, Thomas J, Millam, K. C., Mast, A. R., Paterson, T. B., Theim, T. J., Hipp, A. L., … Sytsma, K. J. (2009). Origin, adaptive radiation and diversification of the Hawaiian lobeliads (Asterales: Campanulaceae). Proceedings of the Royal Society B: Biological Sciences, 276, 407–416. 10.1098/rspb.2008.1204

43. Grant, P. R., & Grant, B. R. (2007). How and why species multiply: The radiation of Darwin’s finches. Princeton: Princeton University Press.

44. Groom, S. V. C., Stevens, M. I., & Schwarz, M. P. (2013). Diversification of Fijian halictine bees: Insights into a recent island radiation. Molecular Phylogenetics and Evolution, 68(3), 582–594. 10.1016/j.ympev.2013.04.015

45. Hamilton, T. H., & Rubinoff, I. (1963). Isolation, endemism, and multiplication of species in the Darwin finches. Evolution, 17(4), 388–403. 10.1111/j.1558-5646.1963.tb03296.x

46. Hembry, D. H., Bennett, G., Bess, E., Cooper, I., Jordan, S., Liebherr, J., … O’Grady, P. M. (2021). Insect Radiations on Islands : Biogeographic pattern and evolutionary process in Hawaiian insects. The Quarterly Review of Biology, 96(4), 247–296. 10.1086/717787

47. Hernández-Hernández, T. (2019). Evolutionary rates and adaptive radiations. Biology & Philosophy, 34(4), 41. 10.1007/s10539-019-9694-y

48. Hiller, A. E., Koo, M. S., Goodman, K. R., Shaw, K. L., O’Grady, P. M., & Gillespie, R. G. (2019). Niche conservatism predominates in adaptive radiation: Comparing the diversification of Hawaiian arthropods using ecological niche modelling. Biological Journal of the Linnean Society, 127(2), 479–492. 10.1093/biolinnean/blz023

49. Hortal, J., Triantis, K. A., Meiri, S., Thébault, E., & Sfenthourakis, S. (2009). Island species richness increases with habitat diversity. The American Naturalist, 174(6), E205–E217. 10.1086/645085

50. Illera, J. C., Rando, J. C., Melo, M., Valente, L., & Stervander, M. (2024). Avian Island Radiations Shed Light on the Dynamics of Adaptive and Nonadaptive Radiation. In C. L. Peichel, D. I. Bolnick, Å. Brännström, U. Dieckmann, & J. S. Rebecca (Éds.), Additional Perspectives on Speciation (p.1–29). Cold Spring Harbor Laboratory Press.

51. Irl, S. D., Harter, D. E., Steinbauer, M. J., Gallego Puyol, D., Fernández-Palacios, J. M., Jentsch, A., & Beierkuhnlein, C. (2015). Climate vs. Topography–spatial patterns of plant species diversity and endemism on a high-elevation island. Journal of Ecology, 103(6), 1621–1633. 10.1111/1365-2745.12463

52. Itow, S. (1995). Phytogeography and ecology of *Scalesia* (Compositae) endemic to the Galápagos Islands. Pacific Science, 49(1), 17–30.

53. Johns, C. A., Toussaint, E. F. A., Breinholt, J. W., & Kawahara, A. Y. (2018). Origin and macroevolution of micro-moths on sunken Hawaiian Islands. Proceedings of the Royal Society B: Biological Sciences, 285, 20181047. 10.1098/rspb.2018.1047

54. Johnson, M. A. (2017). Four new species of *Cyrtandra* (Gesneriaceae) from the South Pacific islands of Fiji. PhytoKeys, 91, 105–124. 10.3897/phytokeys.91.21623

55. Johnson, M. A. (2023). Phylogenetic and functional trait-based community assembly within Pacific Cyrtandra (Gesneriaceae): Evidence for clustering at multiple spatial scales. Ecology and Evolution, 13(5), e10048. 10.1002/ece3.10048

56. Jorgensen, T. H., & Olesen, J. M. (2001). Adaptive radiation of island plants: Evidence from Aeonium (Crassulaceae) of the Canary Islands. Perspectives in Plant Ecology, Evolution and Systematics, 4(1), 29–42. 10.1078/1433-8319-00013

57. Juan, C., Emerson, B. C., Oromı’, P., & Hewitt, G. M. (2000). Colonization and diversification: Towards a phylogeographic synthesis for the Canary Islands. Trends in Ecology & Evolution, 15(3), 104–109.

58. Kalmar, A., & Currie, D. J. (2006). A global model of island biogeography. Global Ecology and Biogeography, 15(1), 72–81. 10.1111/j.1466-822X.2006.00205.x

59. Kim, S.-C., McGowen, M. R., Lubinsky, P., Barber, J. C., Mort, M. E., & Santos-Guerra, A. (2008). Timing and tempo of early and successive adaptive radiations in Macaronesia. PLoS ONE, 3(5), e2139. 10.1371/journal.pone.0002139

60. Kisel, Y., & Barraclough, T. G. (2010). Speciation has a spatial scale that depends on levels of gene flow. The American Naturalist, 175(3), 316–334. 10.1086/650369

61. Kreft, H., Jetz, W., Mutke, J., Kier, G., & Barthlott, W. (2008). Global diversity of island floras from a macroecological perspective. Ecology letters, 11(2), 116–127. 10.1111/j.1461-0248.2007.01129.x

62. Lasky, J. R., Keitt, T. H., Weeks, B. C., & Economo, E. P. (2017). A hierarchical model of whole assemblage island biogeography. Ecography, 40(8), 982–990. 10.1111/ecog.02303

63. Lim, J. Y., & Marshall, C. R. (2017). The true tempo of evolutionary radiation and decline revealed on the Hawaiian archipelago. Nature, 543(7647), 710–713. 10.1038/nature21675

64. Liu, C., Sarnat, E. M., Friedman, N. R., Hita Garcia, F., Darwell, C., Booher, D., … Economo, E. P. (2020). Colonize, radiate, decline: Unraveling the dynamics of island community assembly with Fijian trap-jaw ants. Evolution, 74(6), 1082–1097. 10.1111/evo.13983

65. Losos, J. B., & Ricklefs, R. E. (2009). Adaptation and diversification on islands. Nature, 457(7231), 830–836. 10.1038/nature07893

66. Losos, J. B., & Schluter, D. (2000). Analysis of an evolutionary species–area relationship. Nature, 408(6814), 847–850. 10.1038/35048558

67. MacArthur, R. H., & Wilson, E. O. (1963). An equilibrium theory of insular zoogeography. Evolution, 17(4), 373. 10.2307/2407089

68. MacArthur, R. H., & Wilson, E. O. (1967). The theory of island biogeography. Princeton: Princeton university press.

69. Machado, A., Rodríguez-Expósito, E., López, M., & Hernández, M. (2017). Phylogenetic analysis of the genus *Laparocerus*, with comments on colonisation and diversification in Macaronesia (Coleoptera, Curculionidae, Entiminae). ZooKeys, 651, 1–77. 10.3897/zookeys.651.10097

70. Matthews, T. J., Guilhaumon, F., Triantis, K. A., Borregaard, M. K., & Whittaker, R. J. (2016). On the form of species–area relationships in habitat islands and true islands. Global Ecology and Biogeography, 25(7), 847–858.

71. Matthews, T. J., Rigal, F., Triantis, K. A., & Whittaker, R. J. (2019). A global model of island species–area relationships. *Proceedings of the National Academy of Sciences*, USA, 116(25), 12337–12342. 10.1073/pnas.1818190116

72. Matthews, T. J., Wayman, J. P., Whittaker, R. J., Cardoso, P., Hume, J. P., Sayol, F., … Rigal, F. (2023). A global analysis of avian island diversity–area relationships in the Anthropocene. Ecology Letters, 26(6), 965–982. 10.1111/ele.14203

73. Messerschmid, T. F. E., Abrahamczyk, S., Bañares-Baudet, Á., Brilhante, M. A., Eggli, U., Hühn, P., … Kadereit, G. (2023). Inter- and intra-island speciation and their morphological and ecological correlates in *Aeonium* (Crassulaceae), a species-rich Macaronesian radiation. Annals of Botany, 131(4), 697–721. 10.1093/aob/mcad033

74. Naciri, Y., & Linder, H. P. (2020). The genetics of evolutionary radiations. Biological Reviews, 95(4), 1055–1072. 10.1111/brv.12598

75. Nakagawa, S., & Schielzeth, H. (2013). A general and simple method for obtaining R^2^ from generalized linear mixed-effects models. Methods in ecology and evolution, 4(2), 133–142. 10.1111/j.2041-210x.2012.00261.x

76. Norder, S. J., Proios, K., Whittaker, R. J., Alonso, M. R., Borges, P. A. V., Borregaard, M. K., … Rijsdijk, K. F. (2018). Beyond the Last Glacial Maximum: Island endemism is best explained by long-lasting archipelago configurations. Global Ecology and Biogeography, 28(2), 184–197. 10.1111/geb.12835

77. O’Grady, P., & DeSalle, R. (2018). Hawaiian *Drosophila* as an Evolutionary Model Clade: Days of Future Past. BioEssays, 40(5), 1700246. 10.1002/bies.201700246

78. O’Hara, R., & Kotze, J. (2010). Do not log-transform count data. Nature Precedings, 1–1. 10.1038/npre.2010.4136.1

79. Otto, R., Whittaker, R. J., von Gaisberg, M., Stierstorfer, C., Naranjo-Cigala, A., Steinbauer, M. J., … Fernández-Palacios, J. M. (2016). Transferring and implementing the general dynamic model of oceanic island biogeography at the scale of island fragments: The roles of geological age and topography in plant diversification in the Canaries. Journal of Biogeography, 43(5), 911–922. 10.1111/jbi.12684

80. Parent, C. E. (2012). Biogeographical and ecological determinants of land snail diversification on Islands. American Malacological Bulletin, 30(1), 207–215. 10.4003/006.030.0118

81. Parent, C. E., Caccone, A., & Petren, K. (2008). Colonization and diversification of Galápagos terrestrial fauna: A phylogenetic and biogeographical synthesis. Philosophical Transactions of the Royal Society B: Biological Sciences, 363(1508), 3347–3361. 10.1098/rstb.2008.0118

82. Parent, C. E., & Crespi, B. J. (2006). Sequential colonization and diversification of Galápagos endemic land snail genus *Bulimulus* (Gastropoda, Stylommatophora). Evolution, 60(11), 2311–2328. 10.1111/j.0014-3820.2006.tb01867.x

83. Paulay, G. (1994). Biodiversity on oceanic islands: Its origin and extinction. American zoologist, 34(1), 134–144.

84. Price, J. P., & Wagner, W. L. (2011). A phylogenetic basis for species-area relationships among three Pacific Island floras. American Journal of Botany, 98(3), 449–459. 10.3732/ajb.1000388

85. Price, J. P., & Wagner, W. L. (2018). Origins of the Hawaiian flora: Phylogenies and biogeography reveal patterns of long-distance dispersal. Journal of Systematics and Evolution, 56(6), 600–620. 10.1111/jse.12465

86. R Core Team. (2023). R: A Language and Environment for Statistical Computing. R Foundation for Statistical Computing, Vienna, Austria. <https://www.R-project.org/>.

87. Roell, Y. E., Phillips, J. G., & Parent, C. E. (2021). Effect of topographic complexity on species richness in the Galápagos Islands. Journal of Biogeography, 48(10), 2645–2655. 10.1111/jbi.14230

88. Román-Palacios, C., & Wiens, J. J. (2018). The Tortoise and the Finch: Testing for island effects on diversification using two iconic Galápagos radiations. Journal of Biogeography, 45(8), 1701–1712. 10.1111/jbi.13366

89. Rudbeck, A. V., Sun, M., Tietje, M., Gallagher, R. V., Govaerts, R., Smith, S. A., … Eiserhardt, W. L. (2022). The Darwinian shortfall in plants: Phylogenetic knowledge is driven by range size. Ecography, 2022(8), e06142.

90. Sakai, A. K., Weller, S. G., Wagner, W. L., Nepokroeff, M., & Culley, T. M. (2006). Adaptive radiation and evolution of breeding systems in *Schiedea* (Caryophyllaceae), an endemic Hawaiian genus. Annals of the Missouri Botanical Garden, 93(1), 49–63.

91. Sanmartín, I., Van Der Mark, P., & Ronquist, F. (2008). Inferring dispersal : A Bayesian approach to phylogeny-based island biogeography, with special reference to the Canary Islands. Journal of Biogeography, 35(3), 428–449. 10.1111/j.1365-2699.2008.01885.x

92. Sarnat, E. M., & Economo, E. P. (2012). The ants of Fiji. Univ of California Press.

93. Schenk, J. J. (2021). The Next Generation of Adaptive Radiation Studies in Plants. International Journal of Plant Sciences, 182(4), 245–262. 10.1086/713445

94. Schluter, D. (2000). The ecology of adaptive radiation. Oxford: Oxford University Press.

95. Shaw, K. L., & Gillespie, R. G. (2016). Comparative phylogeography of oceanic archipelagos : Hotspots for inferences of evolutionary process. Proceedings of the National Academy of Sciences, 113(29), 7986–7993. 10.1073/pnas.1601078113

96. Stuessy, T. F., Jakubowsky, G., Gómez, R. S., Pfosser, M., Schlüter, P. M., Fer, T., … Kato, H. (2006). Anagenetic evolution in island plants. Journal of Biogeography, 33(7), 1259–1265. 10.1111/j.1365-2699.2006.01504.x

97. Tao, R., Sack, L., & Rosindell, J. (2021). Biogeographic Drivers of Evolutionary Radiations. Frontiers in Ecology and Evolution, 9, 644328. 10.3389/fevo.2021.644328

98. Taylor, R.J. (1987). The geometry of colonization: 1. Islands. Oikos, 225–231. 10.2307/3565859

99. Triantis, K. A., Guilhaumon, F., & Whittaker, R. J. (2012). The island species-area relationship : Biology and statistics: The island species-area relationship. Journal of Biogeography, 39(2), 215–231. 10.1111/j.1365-2699.2011.02652.x

100. Triantis, K. A., Whittaker, R. J., Fernández-Palacios, J. M., & Geist, D. J. (2016). Oceanic archipelagos : A perspective on the geodynamics and biogeography of the World’s smallest biotic provinces. Frontiers of Biogeography.

101. Vitales, D., Pellicer Moscardó, J., Vallès Xirau, J., & Garnatje i Roca, T. (2015). Molecular insights into the diversification of *Cheirolophus* (Asteraceae) in Macaronesia. In Recent Advances in Pharmaceutical Sciences V (p. 85–100). Research Signpost.

102. Voeten, C. (2023). Buildmer : Stepwise Elimination and Term Reordering for Mixed-Effects Regression. R package version 2.8, <https://CRAN.R-project.org/package=buildmer>.

103. Wagner, C. E., Harmon, L. J., & Seehausen, O. (2014). Cichlid species-area relationships are shaped by adaptive radiations that scale with area. Ecology Letters, 17(5), 583–592. 10.1111/ele.12260

104. Warren, B. H., Simberloff, D., Ricklefs, R. E., Aguilée, R., Condamine, F. L., Gravel, D., … Thébaud, C. (2015). Islands as model systems in ecology and evolution : Prospects fifty years after MacArthur-Wilson. Ecology Letters, 18(2), 200–217. 10.1111/ele.12398

105. Weigelt, P., & Kreft, H. (2013). Quantifying island isolation—Insights from global patterns of insular plant species richness. Ecography, 36(4), 417–429. 10.1111/j.1600-0587.2012.07669.x

106. Weigelt, P., Steinbauer, M. J., Cabral, J. S., & Kreft, H. (2016). Late Quaternary climate change shapes island biodiversity. Nature, 532(7597), 99–102. 10.1038/nature17443

107. White, O. W., Reyes-Betancort, J. A., Chapman, M. A., & Carine, M. A. (2020). Geographical isolation, habitat shifts and hybridisation in the diversification of the Macaronesian endemic genus *Argyranthemum* (Asteraceae). New Phytologist, 228(6), 1953–1971.

108. Whittaker, R. J., Fernández-Palacios, J. M., & Matthews, T. J. (2023). Island Biogeography : Geo-environmental Dynamics, Ecology, Evolution, Human Impact, and Conservation. Oxford University Press.

109. Whittaker, R. J., Triantis, K. A., & Ladle, R. J. (2008). A general dynamic theory of oceanic island biogeography. Journal of Biogeography, 35(6), 977–994. 10.1111/j.1365-2699.2008.01892.x

110. Yang, Y., Morden, C. W., Sporck-Koehler, M. J., Sack, L., Wagner, W. L., & Berry, P. E. (2018). Repeated range expansion and niche shift in a volcanic hotspot archipelago : Radiation of C 4 Hawaiian *Euphorbia* subgenus *Chamaesyce* (Euphorbiaceae). Ecology and Evolution, 8(16), 8523–8536. 10.1002/ece3.4354

111. Yu, F., Wang, T., Groen, T. A., Skidmore, A. K., Yang, X., Geng, Y., & Ma, K. (2015). Multi-scale comparison of topographic complexity indices in relation to plant species richness. Ecological complexity, 22, 93–101.

112. Zuur, A. F., Ieno, E. N., Walker, N. J., Saveliev, A. A., & Smith, G. M. (2009). Mixed effects models and extensions in ecology with R (Vol. 574). Springer.

