## Supplementary Information for "The biogeography of evolutionary radiations on oceanic archipelagos"

##### **Appendix 1**

**Island selection for Hawaii**

The number of Hawaiian Islands historically included in island biogeography analyses have varied greatly between studies (from 5 to 18 islands; Price 2004, Price & Wagner 2011). After a careful review of the literature, we made our own selection by considering biological and statistical matters, as detailed hereafter. First, we remove Kure, Midway, and Pearl & Hermes, because of their islet nature (see Whittaker et al. 2008) and because we consider them too degraded/advanced in their island ontogeny. Then, we excluded Kaula Rock and Kaho‘olawe, due to their highly degraded biota resulting from military explosives testing and alien ungulates presence (King 1973, Whittaker et al. 2008 and reference herein). Finally, we choose to remove Lisianski and Laysan because they are small, far, and old relative the main archipelago and then are strong outliers in most of the geo-environmental factors. Our final set for the main analyses therefore comprises nine islands: Necker, Nihoa, Ni‘ihau, Kauaʻi, Oʻahu, Molokaʻi, Lāna’i, Maui and Hawai’i. Among the 53 ROAs (Evolutionary radiations on oceanic archipelagos) retrieved for Hawaii, 33 ROAs were on 6 islands, 12 on 7 islands, 7 on 8 islands, and 1 on 9 islands.

**Island characteristics**

***Area and environmental heterogeneity***

We used the database of Global Administrative Areas (GADM http://www.gadm.org/version1) to obtain high-resolution present coastlines for all islands included in our dataset. We then gathered information for island area, environmental heterogeneity, geological age and intra-archipelagic island isolation as potential determinants of Island SROA. Environmental heterogeneity was assessed using two distinct variables: (1) the maximum elevation – used here as a proxy of the climatic gradient and habitat diversity, and; (2) the variation in the rate of elevational change over the horizontal surface – quantified here as the standard deviation of slope (hereafter SDS) – which is a measure of the topographic complexity of an island. The maximum elevation and SDS were calculated based on the 30-m digital elevation model (DEM) Advanced Land Observing Satellite (ALOS) from the OpenTopography Portal (<https://opentopography.org>).

***Neighbor Index (NI) calculation***

The Neighbor Index (NI) (Kalmar & Currie 2006) postulated that the importance of an island as a potential source of colonization to the focal island is proportional to its area (A) and inversely proportional to the distance (*D*) that separates it from the focal island. The general formula of NI for a given island is written as follow:

$$NI=\sum_{i=1}^{n-1} \frac{A_{i}^{a}}{D_{i}^{b}}$$

where *n* is the number of islands in the archipelago, and *a* and *b* are empirical constants. Kalmar & Curie (2006) suggested estimating *a* and *b* iteratively for each dataset until they maximize the correlation with RSI. Because we have few data points per ROA, we did not follow this approach to avoid spurious values for *a* and *b*. Instead, we identified three significant pairs of *a* and *b,* and calculated NI for each ROA using the pair that maximizes the correlation with RSI. The three pairs were: (1) *a* = 0 and *b* = - 1, where NI is simply the sum of the distance (no effect of area); (2) *a* = 1 and *b* = 1, where NI is proportional to the area, and inversely proportional the distance and (3) *a* = 1 and *b* = 2, where NI is proportional to their area, and inversely proportional to the square of their distance. Each of the three NI calculations was done specifically for each ROA, only accounting for islands where the ROA occurs. The results showed that the best configuration for measuring inter-island isolation was the one that only took distance into account in its calculation for 65% of ROAs, while 28% of ROAs showed the best correlation with area and distance squared, and only 7% with area and distance.

**Finding the best random structure for GLMM**

The best random effect structures, with all fixed effects considered, were selected using the function buildmer in the R package *buildmer* (Voeten 2023). The function performs a backward stepwise elimination of the random terms while guaranteeing model convergence at the same time, which is a critical issue for models with complex random structures such as ours. The backward stepwise procedure was performed using the Likelihood Ratio Test (LRT) as implemented by default. The selection of the best random structures for the SIE analysis was performed using the *buildglmmTMB* function in the *buildmer* package.

**References**

Ewing, C. (2007). Phylogenetic analysis of the genera of endemic Hawaiian sap beetles (Coleoptera: Nitidulidae) based on morphology with redescription and key to the genera of endemic Hawaiian Nitidulidae. *Zootaxa*, *1427*(1), 1–36. <https://doi.org/10.11646/zootaxa.1427.1.1>

Gero, P. D. (2015). *Radiation of the bark louse genus* Kilauella *across the Hawaiian Islands* [PhD Thesis]. University of Illinois.

Goodman, K. R., Evenhuis, N. L., Bartošová-Sojková, P., & O’Grady, P. M. (2014). Diversification in Hawaiian long-legged flies (Diptera: Dolichopodidae: *Campsicnemus*): Biogeographic isolation and ecological adaptation. *Molecular Phylogenetics and Evolution*, *81*, 232–241.

<https://doi.org/10.1016/j.ympev.2014.07.015>

Hembry, D. H., Bennett, G., Bess, E., Cooper, I., Jordan, S., Liebherr, J., Magnacca, K. N., Percy, D. M., Polhemus, D. A., Rubinoff, D., Shaw, K. L., & O’Grady, P. M. (2021). Insect Radiations on Islands: Biogeographic Pattern and Evolutionary Process in Hawaiian Insects. *The Quarterly Review of Biology*, *96*(4), 247–296. <https://doi.org/10.1086/717787>

Holland, B. S., & Cowie, R. H. (2009). Land Snail Models in Island Biogeography: A Tale of Two Snails^*^. *American Malacological Bulletin*, *27*(1–2), 59–68. <https://doi.org/10.4003/006.027.0205>

Kalmar, A., & Currie, D. J. (2006). A global model of island biogeography. *Global Ecology and Biogeography*, *15*(1), 72–81. <https://doi.org/10.1111/j.1466-822X.2006.00205.x>

King, B. (1973). Conservation status of birds of central pacific islands. *The Wilson Bulletin*, *85*(1), 89–103.

O’Grady, P., & DeSalle, R. (2018). Hawaiian *Drosophila* as an Evolutionary Model Clade: Days of Future Past. *BioEssays*, *40*(5), 1700246. <https://doi.org/10.1002/bies.201700246>

Price, J. P. (2004). Floristic biogeography of the Hawaiian Islands: Influences of area, environment and paleogeography: Floristic biogeography of the Hawaiian Islands. *Journal of Biogeography*, *31*(3), 487–500. <https://doi.org/10.1046/j.0305-0270.2003.00990.x>

Price, J. P., & Wagner, W. L. (2011). A phylogenetic basis for species-area relationships among three Pacific Island floras. *American Journal of Botany*, *98*(3), 449–459. <https://doi.org/10.3732/ajb.1000388>

Smith, A. C. (1991). *Flora vitiensis nova. A new flora of Fiji* (Vol. 5). National Tropical Botanical Garden.

Voeten, C. (2023). *Buildmer:* Stepwise Elimination and Term Reordering for Mixed-Effects Regression. R package version 2.8, <https://CRAN.R-project.org/package=buildmer>.

Whittaker, R. J., Triantis, K. A., & Ladle, R. J. (2008). A general dynamic theory of oceanic island biogeography: A general dynamic theory of oceanic island biogeography. *Journal of Biogeography*, *35*(6), 977–994. <https://doi.org/10.1111/j.1365-2699.2008.01892.x>

##### **Table S1.** List of the 20 oceanic archipelagos fitting all our criteria (i.e. fully oceanic with at least six main islands) and results of our literature research for ROA description. FP=French Polynesia; MAC=Macaronesia.

| **Archipelago name** | **ROA presence**  (10 species or more) |
| --- | --- |
| Austral (FP) | Yes |
| Azores (MAC) | Yes |
| Canary Islands (MAC) | Yes |
| Cabo Verde (MAC) | Yes |
| Cook | No |
| Faroe | No |
| Fiji | Yes |
| Galápagos | Yes |
| Gambier (FP) | No |
| Hawaii | Yes |
| Kermadec | No |
| Lesser Antilles | Yes |
| Line islands | No |
| Mariana | No |
| Marquesas (FP) | Yes |
| Ogasawara (Bonin) | Yes |
| Samoa | No |
| Society (FP) | Yes |
| Tonga | No |
| Vanuatu | Yes |
| Aleutian Islands | No |
| Kuril Islands | No |
| South Sandwich Islands | No |
| Californian Channel Islands | No |
| Aeolian Islands | No |
| Palau | No |
| Tuamotu Archipelago | No |
| Lakshadweep | No |

##### **Table S2.** List of the ROAs with at least 10 species and occurring on our four selected oceanic archipelagos but removed from our final dataset for specific reasons detailed below.

| **Archipelago** | **ROA name** | **Reason for removal** |
| --- | --- | --- |
| Hawaii | *Tetragnatha*  (Web Builders) | Distributed on only five islands |
|  | *Orsonwelles* | Distributed on only five islands |
|  | *Lysimachia* | Distributed on only five islands |
|  | *Auricullellinae* | Distributed on only five islands |
|  | *Hyposmocoma* | Too many undescribed species: at least 600 species are known, but hundreds of them are yet to be formerly described (Hembry et al. 2021) |
|  | *Kilauella* | Too many undescribed species: over 200 species are estimated based on museum collections, but only seven of them have been published (Gero 2015, Kevin Johnson personal communication) |
|  | Sap beetles | Too many undescribed species (see Ewing 2007) |
|  | Succineidae | Succineidae land snails have colonized the Hawaiian Islands twice according to molecular data (Holland & Cowie 2009), but the species repartition between those two independent colonizations can't be assessed properly. |
|  | *Campsicnemus* | Too many undescribed species: Goodman et al. (2014) estimated that 60 species await description, and most of them have yet to be published to date. |
|  | *Drosophila* | Too many undescribed species: hundreds of species are still awaiting description (O’Grady & DeSalle 2018) |
| Canary Islands | *Calathus* | Distributed on only five islands |
|  | *Cheirolophus* | Distributed on only five islands |
|  | *Pericallis* | Distributed on only five islands |
| Fiji | *Copelatus* | Distributed on only five islands |
|  | *Fluviopupa* | Distributed on only two islands |
|  | *Phyllanthus* | The only published reference of *Phyllanthus* data distribution is Smith (1991) book "Flora Vitiensis Nova" (David H. Hembry personal communication), which lacks information/precision on the matter. |
| Society | *Rhyncogonus* | Distributed on only five islands |

**Table S3**. List of the ROAs selected for the study with their associated characteristics.

##### **Table S4.** Islands characteristics for the four oceanic archipelagos selected in our study. SDS correspond to the standard deviation of the slope (see main text). The three _NI columns correspond to the three Neighbor Index calculations (D_NI = only distances, AD_NI = area & distances, AD²_NI = area & distances², see main text and **Appendix 1**. Note: These values of NI are indicative since, for a given ROA, they have been calculated only with the islands on which the species constituent of the ROA occurs.

**Table S5.** Summary of the results of the univariate GLMs between Island S_ROA_ and each geo-environmental factor separately for the five retrieved archipelagos with less than five ROAs each, namely Society, Marquesas, Austral Islands, Azores and Cabo Verde. For each archipelago, the total number of ROAs retrieved is given in parentheses. For each factor and for each archipelago, the number of significant relationships between the factor and Island S_ROA_ is given before and after the FDR correction (separated by a vertical bar). Specifically for age + age^2^, only the number of radiations having a hump-shaped relationship with age that met our three criteria is given.

|  | Archipelagos | | | | |
| --- | --- | --- | --- | --- | --- |
| Factors | Austral Islands (1) | Azores (1) | Cabo Verde (1) | Marquesas (3) | Society (4) |
| Area | 0 | 0 | 0 | 0 | 3\|1 |
| Age | 0 | 0 | 0 | 0 | 3\|2 |
| Age+Age^2^ | 0 | 0 | 0 | 0 | 0 |
| Elevation | 1\|1 | 0 | 0 | 0 | 3\|2 |
| SDS | 1\|1 | 0 | 0 | 0 | 3\|2 |
| Inter-island isolation | 1\|1 | 0 | 0 | 0 | 4\|1 |

**Table S6.** The best random structure and family distribution selected for the GLMMs performed for data including all current geo-environmental factors for our initial model (see Main text) and the model including the subspecies. The best random structures were selected using the function *buildmer* in the R package *buildmer* (Voeten 2023). The function performs a backward stepwise elimination of the random terms while guaranteeing model convergence at the same time. A negative binomial was selected when the GLMM with Poisson showed significant overdispersion.

|  | Best random structure | |  |
| --- | --- | --- | --- |
|  | Random slope | Random intercept | Family distribution |
| Initial model |  |  |  |
| Hawaii | elevation, inter-island isolation | Radiations, Islands | Poisson |
| Canary Islands | SDS, age | Radiations | Poisson |
| Galápagos | SDS | Radiations | Poisson |
| Fiji | - | Radiations | Poisson |
| Plants Hawaii | Elevation, age^2^ | Radiations | Poisson |
| Plants Canary Islands |  | Radiations | Poisson |
| Invertebrates Hawaii | SDS, inter-island isolation | Radiations, Islands | Poisson |
| Invertebrates Canary Islands | SDS, age | Radiations | Poisson |
| Subspecies models |  |  |  |
| Hawaii | SDS, inter-island isolation | Radiations, Islands | Poisson |
| Canary | SDS, elevation | Radiations | Poisson |
| Galápagos | SDS | Radiations | Poisson |
| Fiji | - | Radiations | Poisson |
| Plants Hawaii | Elevation, age^2^ | Radiations | Poisson |
| Plants Canary Islands |  | Radiations | Poisson |
| Invertebrates Hawaii | SDS, elevation, inter-island isolation | Radiations, Islands | Poisson |
| Invertebrates Canary Islands | SDS | Radiations | Poisson |

**Table S7.** Results of the ANOVAs testing whether the strength of the effect of the geo-environmental factors on the Island S_ROA_ can be explained by archipelagos, taxa as well as radiation size (log_10_-tranformed) and proportion of SIE. The analyses were performed with subspecies. The effect of a given geo-environmental factor on the Island S_ROA_ of a given ROA is quantified by its standardized β coefficient for the factors area, age, elevation, SDS and inter-island isolation, while the effect of age + age^2^ is coded as binary variable (See main text). For the age + age^2^ model, the model for then fitted with a binomial. The R^2^ is given for each model and the *P*-value is provided for each predictor remaining in the model after backward elimination procedure. Analyses were initially performed for all radiations without testing the interaction between Archipelago and Taxa because of a too unbalanced design. The models with the interaction term were performed for a subset of the data including only the radiations of Hawaii and the Canary Islands and for plants and invertebrates. Note that archipelago and taxa were kept in the model when their interaction was significant even if their independent effect was not. Significant effects are marked in bold.

|  |  | | Predictors | | | | | | | |  |
| --- | --- | --- | --- | --- | --- | --- | --- | --- | --- | --- | --- |
|  | Factors | | Archipelago | | Taxa | A x T | Size | | Proportion of SIE | | R^2^ |
| All radiations |  | |  | |  |  |  | |  | |  |
| Standardized β coefficient | Area | | **<0.001** | | **0.015** | - | **<0.001** | | **0.002** | | 0.528 |
|  | Age | | **0.010** | |  | - |  | |  | | 0.114 |
|  | Age + age2 | | **0.003** | |  | - |  | | **0.047** | | 0.325 |
|  | Elevation | | **0.001** | |  | **-** | **<0.001** | | **0.009** | | 0.382 |
|  | SDS | | **<0.001** | | **<0.001** | - |  | |  | | 0.532 |
|  | Inter-island isolation | |  | |  | - |  | |  | | - |
| Subset | |  | |  | |  | |  | |  | |
| Standardized β coefficient | Area | |  | | **0.022** |  | **<0.001** | | **0.019** | | 0.208 |
|  | Age | | 0.711 | | 0.941 | **<0.001** |  | | **0.029** | | 0.144 |
|  | Age + age2 | | **0.002** | |  |  |  | | **0.049** | | 0.146 |
|  | Elevation | | **0.018** | | 0.928 | **0.001** | **<0.001** | |  | | 0.297 |
|  | SDS | | **0.003** | | **<0.001** |  |  | |  | | 0.274 |
|  | Inter-island isolation | | **0.002** | |  |  |  | |  | | 0.11 |

**Table S8.** Results of the Generalized Linear Mixed Models (GLMMs) analyses testing the effect of the geo-environmental factors on S_ROA_ variation simultaneously for all the radiations in Hawaii and the Canary Islands and for their respective plants and invertebrates subsets. The analyses were performed with subspecies. The number of ROAs involved in each GLMM is given in parentheses with subset’s name. Model selection was performed using AICc. Only the best-fit models (∆AICc < 2) are presented, with their coefficient of the factors included in the model, the degree of freedom (df), the AICc, ∆AICc and *w*AICc. The significance of each factor was estimated using an analysis of variance. Significant factors are marked in bold. It should be noted that the calculation of *P*-values for fixed effects in GLMMs remains controversial. Further details including the best random structure, and family distribution are given in **Table S5**.

|  |  | Geo-environmental factors | | | | | |  |  |  |  |
| --- | --- | --- | --- | --- | --- | --- | --- | --- | --- | --- | --- |
|  | Subsets | Area | Age | Age^2^ | Elevation | SDS | Inter-island isolation | df | AICc | ΔΑΙCc | *w*AICc |
| Hawaii (53) | | **0.500** |  |  | **0.223** | **0.369** |  | 11 | 1966.309 | 0 | 0.310 |
|  |  | **0.494** |  |  | **0.221** | **0.366** | -0.021 | 12 | 1968.043 | 1.734 | 0.130 |
| Canary Islands (29) | |  | **0.108** | **-0.124** | **0.256** |  | **-0.290** | 11 | 961.873 | 0.000 | 0.497 |
| Galapagos (7) | |  |  |  | **0.339** | **-0.294** | -0.104 | 7 | 334.831 | 0.000 | 0.080 |
|  |  | **0.515** | 0.050 | **-0.129** |  |  |  | 7 | 335.122 | 0.292 | 0.069 |
|  |  |  |  |  | **0.361** | **-0.312** |  | 6 | 335.192 | 0.361 | 0.067 |
|  |  |  | 0.056 | **-0.106** | **0.416** | -0.275 |  | 8 | 335.583 | 0.752 | 0.055 |
|  |  | 0.309 | 0.074 | **-0.124** | 0.209 |  |  | 8 | 335.655 | 0.824 | 0.053 |
|  |  |  |  |  | **0.352** |  | -0.108 | 6 | 335.761 | 0.931 | 0.050 |
|  |  |  | 0.078 | **-0.107** | **0.445** |  |  | 7 | 336.166 | 1.335 | 0.041 |
|  |  | **0.453** |  |  |  |  |  | 5 | 336.213 | 1.382 | 0.040 |
|  |  |  |  |  | **0.373** |  |  | 5 | 336.373 | 1.542 | 0.037 |
|  |  | **0.424** |  |  |  |  | -0.092 | 6 | 336.447 | 1.616 | 0.036 |
|  |  |  | 0.050 |  | **0.366** | -0.263 | -0.110 | 8 | 336.670 | 1.839 | 0.032 |
|  |  | 0.248 |  |  | 0.187 |  |  | 6 | 336.816 | 1.985 | 0.030 |
| Fiji (6) | |  | **0.394** | **-0.214** | **0.832** |  |  | 5 | 246.141 | 0 | 0.180 |
|  |  |  | **0.407** | **-0.257** | **0.787** | 0.175 |  | 6 | 246.267 | 0.126 | 0.169 |
|  |  |  | **0.353** | **-0.180** | **0.822** |  | -0.104 | 6 | 247.041 | 0.900 | 0.115 |
|  |  | 0.245 | **0.199** |  | **0.461** |  | -0.138 | 6 | 248.032 | 1.891 | 0.070 |
|  |  |  | **0.375** | **-0.225** | **0.787** | 0.145 | -0.076 | 7 | 248.121 | 1.980 | 0.067 |
| Plants | Hawaii (22) |  | **0.172** | **-0.484** | **0.574** |  |  | 10 | 717.097 | 0 | 0.279 |
|  |  |  | **0.148** | **-0.420** | **0.561** | 0.070 |  | 11 | 717.456 | 0.360 | 0.233 |
|  |  | -0.089 | **0.182** | **-0.504** | **0.657** |  |  | 11 | 718.474 | 1.378 | 0.140 |
|  | Canary Islands (11) |  | -0.029 | **-0.208** | **0.403** |  | **-0.192** | 6 | 317.863 | 0 | 0.213 |
|  |  |  | 0.122 | **-0.214** | **0.527** | 0.157 |  | 6 | 319.002 | 1.140 | 0.121 |
|  |  |  | 0.043 | **-0.199** | **0.425** | 0.097 | -0.152 | 7 | 319.398 | 1.536 | 0.099 |
|  |  |  | 0.025 | **-0.234** | **0.544** |  |  | 5 | 319.521 | 1.659 | 0.093 |
| Invertebrates | Hawaii (30) | **0.517** |  |  | **0.317** | **0.376** |  | 15 | 1185.463 | 0 | 0.255 |
|  |  | **0.794** |  |  |  | **0.436** |  | 14 | 1187.411 | 1.948 | 0.096 |
|  |  | **0.511** |  |  | **0.312** | **0.375** | -0.030 | 16 | 1187.460 | 1.998 | 0.094 |
|  | Canary Islands (18) |  | **0.141** | **-0.092** | **0.167** |  | **-0.352** | 8 | 638.292 | 0 | 0.162 |
|  |  |  | 0.097 | **-0.094** | **0.150** | -0.141 | **-0.387** | 9 | 638.694 | 0.402 | 0.133 |
|  |  | **0.109** | 0.003 | **-0.118** |  |  | **-0.430** | 8 | 638.912 | 0.620 | 0.119 |
|  |  | **0.098** | -0.027 | -**0.117** |  | -0.144 | **-0.458** | 9 | 639.177 | 0.886 | 0.104 |
|  |  |  | **0.160** |  | **0.178** |  | **-0.399** | 7 | 640.164 | 1.873 | 0.064 |

**Table S9.** Results of the post-hoc tests performed on the ANOVAs testing whether the strength of the effect of the geo-environmental factors on the Island S_ROA_ can be explained by archipelagos, taxa as well as radiation size (log_10_-tranformed) and proportion of SIE (See Main text Table 1). Only significant pairwise comparisons are shown.

| All Radiations | Factors | Predictors | Pairwise comparisons | Z-ratio | P |
| --- | --- | --- | --- | --- | --- |
|  | Area |  |  |  |  |
|  |  | Archipelago | Hawaii < Galápagos | -5.856 | <0.001 |
|  |  |  | Canary < Galápagos | -6.309 | <0.001 |
|  |  |  | Galápagos > Fiji | 3.665 | 0.004 |
|  |  | Taxa | Invertebrates > Plants | 2.436 | 0.017 |
|  |  |  | Hawaii < Galápagos | -2.108 | 0.038 |
|  | Age | Archipelago | Hawaii < Fiji | -2.917 | 0.005 |
|  |  |  | Canary < Fiji | -2.461 | 0.016 |
|  | Age+Age2 | Archipelago | Hawaii < Canary | -2.599 | 0.01 |
|  |  |  | Hawaii < Galápagos | -3.867 | <0.001 |
|  | Elevation | Archipelago | Canary < Galápagos | -2.800 | 0.006 |
|  |  |  | Galápagos > Fiji | 2.418 | 0.018 |
|  | SDS | Archipelago | Hawaii > Canary | 4.300 | <0.001 |
|  |  | Taxa | Invertebrates < Vertebrates | -2.494 | 0.015 |
|  |  |  | Invertebrates < Plants | -3.936 | <0.001 |
|  | Inter-island isolation | Archipelago | Hawaii > Canary | 4.555 | <0.001 |
|  |  |  | Canary < Galápagos | -2.226 | 0.029 |
| Subsets | Factors | Predictors | Pairwise comparisons | Z-ratio | P |
|  | Area | Taxa | Invertebrates > Plants | 2.121 | 0.037 |
|  |  |  | Hawaii Invertebrates < Canary Invertebrates | -2.365 | 0.021 |
|  | Age | A x T | Canary Invertebrates > Canary Plants | 2.546 | 0.013 |
|  |  |  | Hawaii Plants > Canary Plants | 2.075 | 0.041 |
|  | Age+Age^2^ | Archipelago | Canary Islands > Hawaii | 2.566 | 0.010 |
|  | Elevation |  |  |  |  |
|  |  | Archipelago | Canary Islands > Hawaii | 3.104 | 0.003 |
|  |  |  | Hawaii Invertebrates < Canary Plants | -2.842 | 0.006 |
|  |  | A x T | Canary Invertebrates < Canary Plants | -3.058 | 0.003 |
|  |  |  | Hawaii Plants < Canary Plants | -4.388 | <0.001 |
|  |  | Archipelago | Canary Islands > Hawaii | 3.047 | 0.003 |
|  | SDS |  |  |  |  |
|  |  | Taxa | Invertebrates < Plants | -4.509 | <0.001 |
|  | Inter-island isolation |  |  |  |  |
|  |  | Archipelago | Canary Islands < Hawaii | -5.111 | <0.001 |
|  |  | Taxa | Invertebrates > Plants | 1.992 | 0.049 |
|  |  |  | Hawaii Invertebrates > Canary Invertebrates | 2.718 | 0.008 |
|  |  |  | Hawaii Invertebrates > Canary Plants | 5.488 | <0.001 |
|  |  | A x T | Canary Invertebrates < Hawaii Plants | -2.220 | 0.029 |
|  |  |  | Canary Invertebrates > Canary Plants | 2.937 | 0.004 |
|  |  |  | Hawaii Plants > Canary Plants | 4.954 | <0.001 |

**Table S10.** Results of the Generalized Linear Mixed Models (GLMMs) analyses testing the effect of the geo-environmental factors on S_ROA_ variation simultaneously for all the radiations in Hawaii and the Canary Islands and for their respective plant and invertebrate subsets. The number of ROAs involved in each GLMM is given in parentheses with subset’s name. Model selection was performed using AICc. Only the best-fit models (∆AICc < 2) are presented, with their coefficient of the factors included in the model, the degree of freedom (df), the AICc, ∆AICc and *w*AICc. The significance of each factor was estimated using an analysis of variance. Significant factors are marked in bold. It should be noted that the calculation of *P*-values for fixed effects in GLMMs remains controversial. Further details including the best random structure, and family distribution are given in **Table S6**.

|  |  | Geo-environmental factors | | | | | |  |  |  |  |
| --- | --- | --- | --- | --- | --- | --- | --- | --- | --- | --- | --- |
|  | Subsets | Area | Age | Age^2^ | Elevation | SDS | inter-island Isolation | df | AICc | ΔΑΙCc | *w*AICc |
| Hawaii (53) | | **0.483** |  |  | 0.201 | **0.359** |  | 11 | 1957.432 | 0 | 0.234 |
|  |  | **0.477** |  |  | 0.172 | **0.345** | -0.074 | 12 | 1958.634 | 1.202 | 0.128 |
|  |  | **0.658** |  |  |  | **0.383** |  | 10 | 1959.051 | 1.619 | 0.104 |
|  |  | **0.613** |  |  |  | **0.356** | -0.108 | 11 | 1959.059 | 1.627 | 0.104 |
|  |  | **0.612** | -0.109 |  |  | **0.407** |  | 11 | 1959.194 | 1.762 | 0.097 |
| Canary Islands (29) | |  | **0.139** | **-0.148** | **0.296** |  | **-0.314** | 11 | 950.562 | 0 | 0.495 |
|  |  | -0.062 | 0.211 | **-0.134** | **0.372** |  | **-0.268** | 12 | 952.258 | 1.696 | 0.212 |
| Galapagos (7) | | **0.515** | 0.050 | **-0.129** |  |  |  | 7 | 335.120 | 0 | 0.078 |
|  |  |  |  |  | **0.361** | **-0.312** |  | 6 | 335.192 | 0.072 | 0.075 |
|  |  |  |  |  | **0.319** | **-0.303** | -0.186 | 7 | 335.536 | 0.416 | 0.063 |
|  |  |  | 0.057 | **-0.106** | **0.416** | -0.275 |  | 8 | 335.577 | 0.457 | 0.062 |
|  |  | 0.308 | 0.075 | **-0.124** | 0.209 |  |  | 8 | 335.650 | 0.530 | 0.060 |
|  |  |  | 0.078 | **-0.107** | **0.445** |  |  | 7 | 336.155 | 1.036 | 0.046 |
|  |  | **0.453** |  |  |  |  |  | 5 | 336.213 | 1.093 | 0.045 |
|  |  |  |  |  | **0.373** |  |  | 5 | 336.373 | 1.253 | 0.042 |
|  |  |  |  |  | **0.330** |  | -0.192 | 6 | 336.498 | 1.378 | 0.039 |
|  |  | 0.248 |  |  | 0.187 |  |  | 6 | 336.816 | 1.697 | 0.033 |
|  |  | **0.403** |  |  |  |  | -0.176 | 6 | 336.839 | 1.720 | 0.033 |
| Fiji (6) | |  | **0.394** | **-0.214** | **0.832** |  |  | 5 | 246.141 | 0 | 0.238 |
|  |  |  | **0.407** | **-0.257** | **0.787** | 0.175 |  | 6 | 246.267 | 0.126 | 0.223 |
| Plants | Hawaii (22) |  | **0.155** | **-0.462** | **0.544** |  |  | 10 | 708.403 | 0 | 0.237 |
|  |  |  | **0.129** | **-0.393** | **0.529** | 0.074 |  | 11 | 708.655 | 0.252 | 0.209 |
|  |  | -0.086 | **0.164** | **-0.482** | **0.625** |  |  | 11 | 709.860 | 1.457 | 0.114 |
|  |  |  | **0.172** | **-0.427** | **0.530** |  | -0.073 | 11 | 710.257 | 1.854 | 0.094 |
|  | Canary Islands (11) |  | 0.023 | **-0.197** | **0.538** |  |  | 6 | 314.193 | 0 | 0.200 |
|  |  |  | 0.109 | **-0.180** | **0.522** | 0.139 |  | 7 | 314.537 | 0.344 | 0.168 |
|  |  | -0.122 | 0.159 | -0.151 | **0.653** |  |  | 7 | 315.774 | 1.581 | 0.091 |
|  |  | **-0.250** | **0.349** |  | **0.827** |  |  | 6 | 315.996 | 1.803 | 0.081 |
| Invertebrates | Hawaii (30) | **0.565** |  |  | 0.304 | **0.390** |  | 11 | 1180.260 | 0 | 0.220 |
|  |  | **0.818** |  |  |  | **0.433** |  | 10 | 1181.589 | 1.329 | 0.113 |
|  |  | **0.553** |  |  | 0.297 | **0.387** | -0.120 | 12 | 1181.816 | 1.556 | 0.101 |
|  | Canary Islands (18) |  | **0.174** | **-0.135** | **0.207** |  | **-0.405** | 11 | 631.344 | 0 | 0.416 |

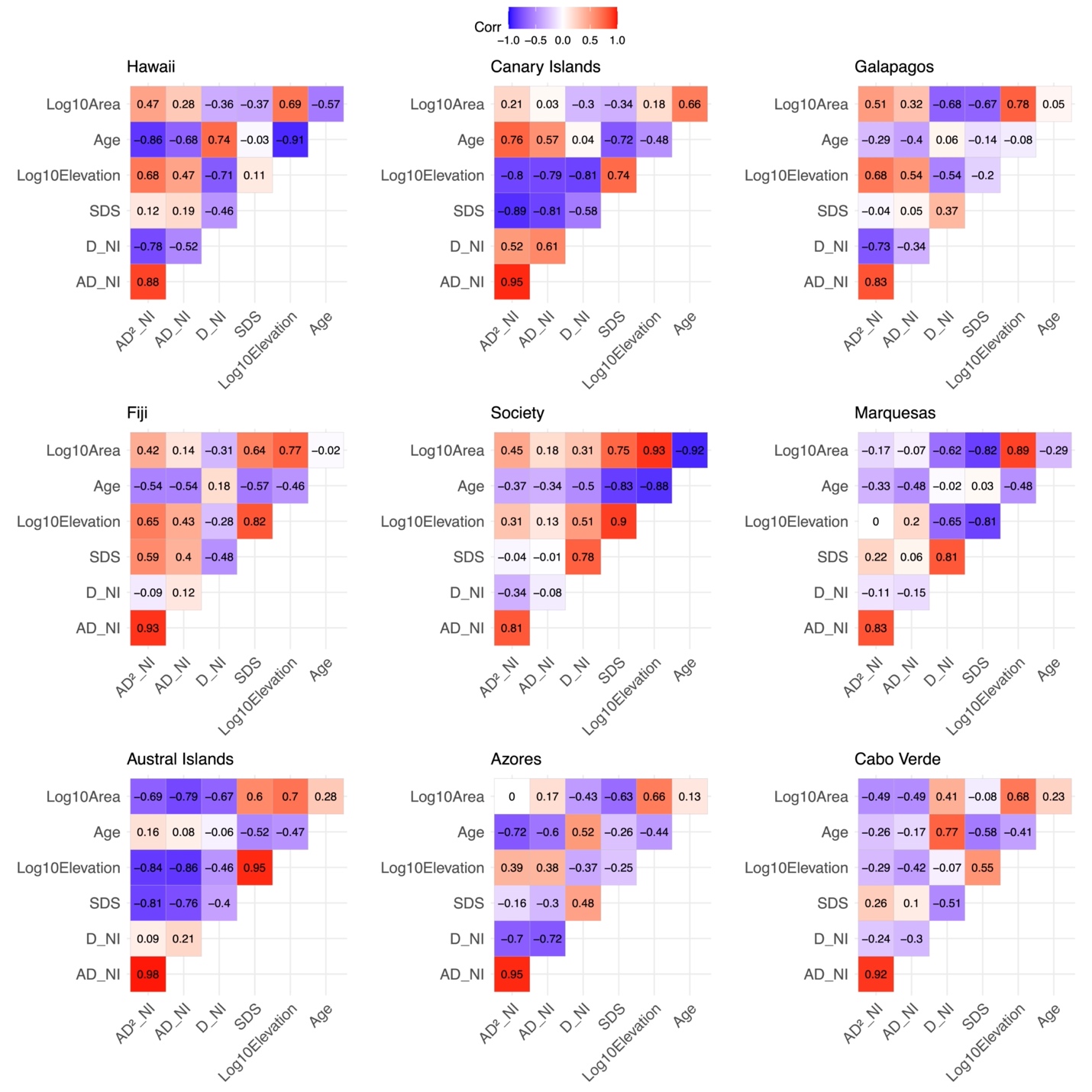

**Figure S1.** Pairwise correlations between the biogeographical factors for each archipelago. SDS: Standard deviation of slopes. D_NI corresponds to the Neighbor Index where NI is simply the sum of the distance (no effect of area); AD_NI where NI is proportional to the area, and inversely proportional the distance and AD^2^_NI where NI is proportional to their area, and inversely proportional to the square of their distance (Values of the geo-environmental factors are given in Table S4).

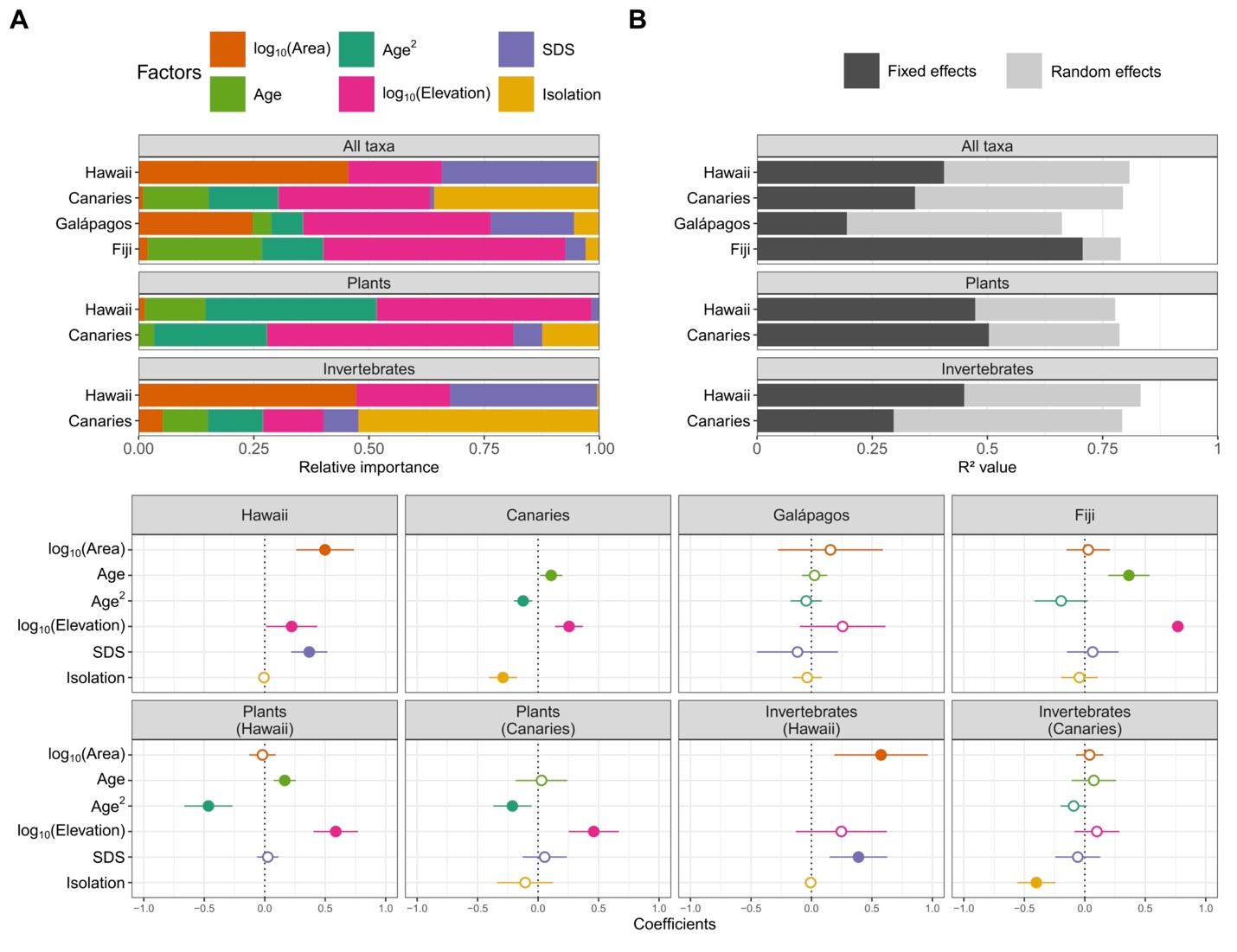

**Figure S2**. Results of the GLMMs performed with the subspecies included. (A) The relative importance of each fixed effect as predictor of the Island S_ROA_ for each for all the radiations in each archipelago and the plant and invertebrate subsets for the Hawaii and Canary Islands. (B) Decomposition of the variance explained by the best set of models (conditional R^2^c) into its components assigned to the fixed effect (marginal R^2^m, dark grey) and random effects effect (R^2^c - R^2^m, light grey) for each archipelago, and the plants and invertebrates subsets (C) Coefficients of geo-environmental factors and their associated confidence intervals extracted from the model averaged over the best models identified for each archipelago and for the subsets of plants and invertebrates. Solid dots indicate a significant effect, while empty dots indicate no effect. Details for the best models are in Table S10.

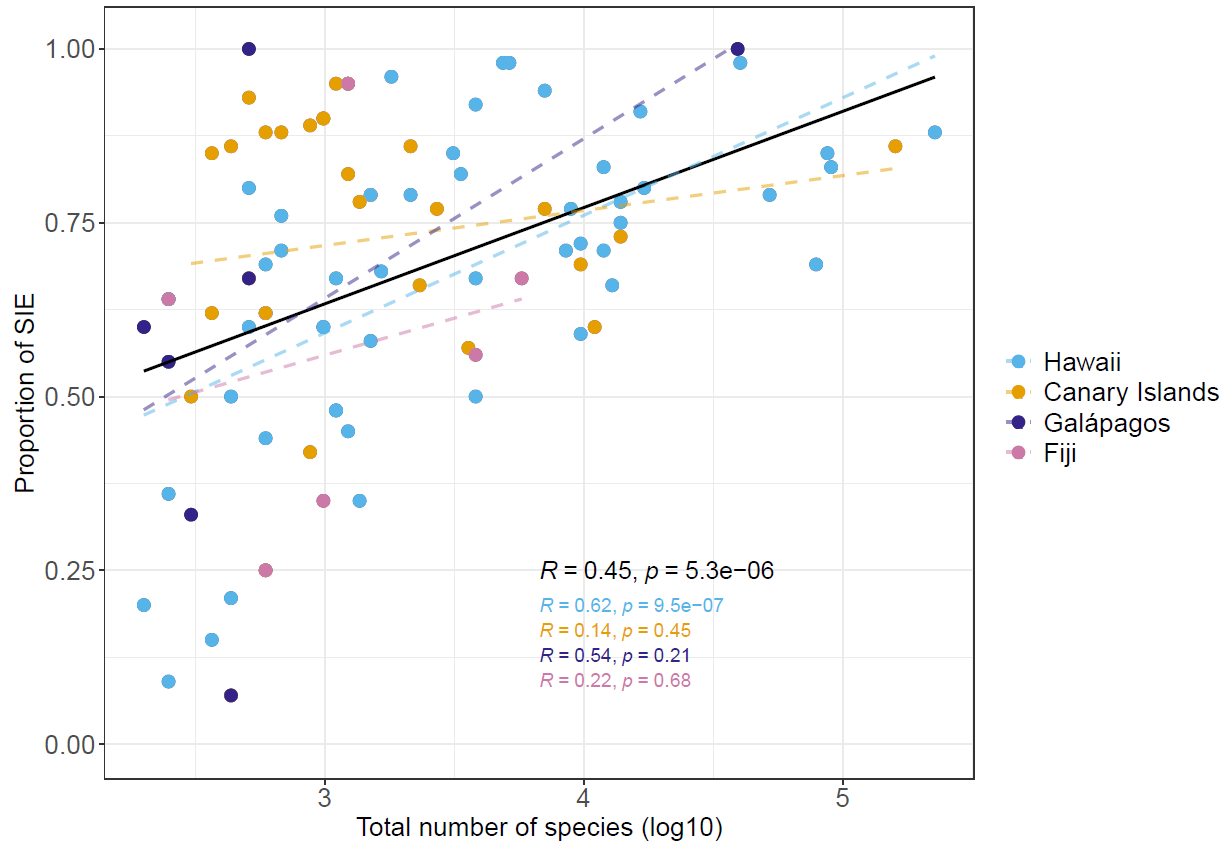

**Figure S3**. Linear relation between ROAs species richness and their percentage of Single Island Endemics (SIE). Black line represents the global linear regression (n=95). Dashed lines represent linear regressions for each archipelago independently.

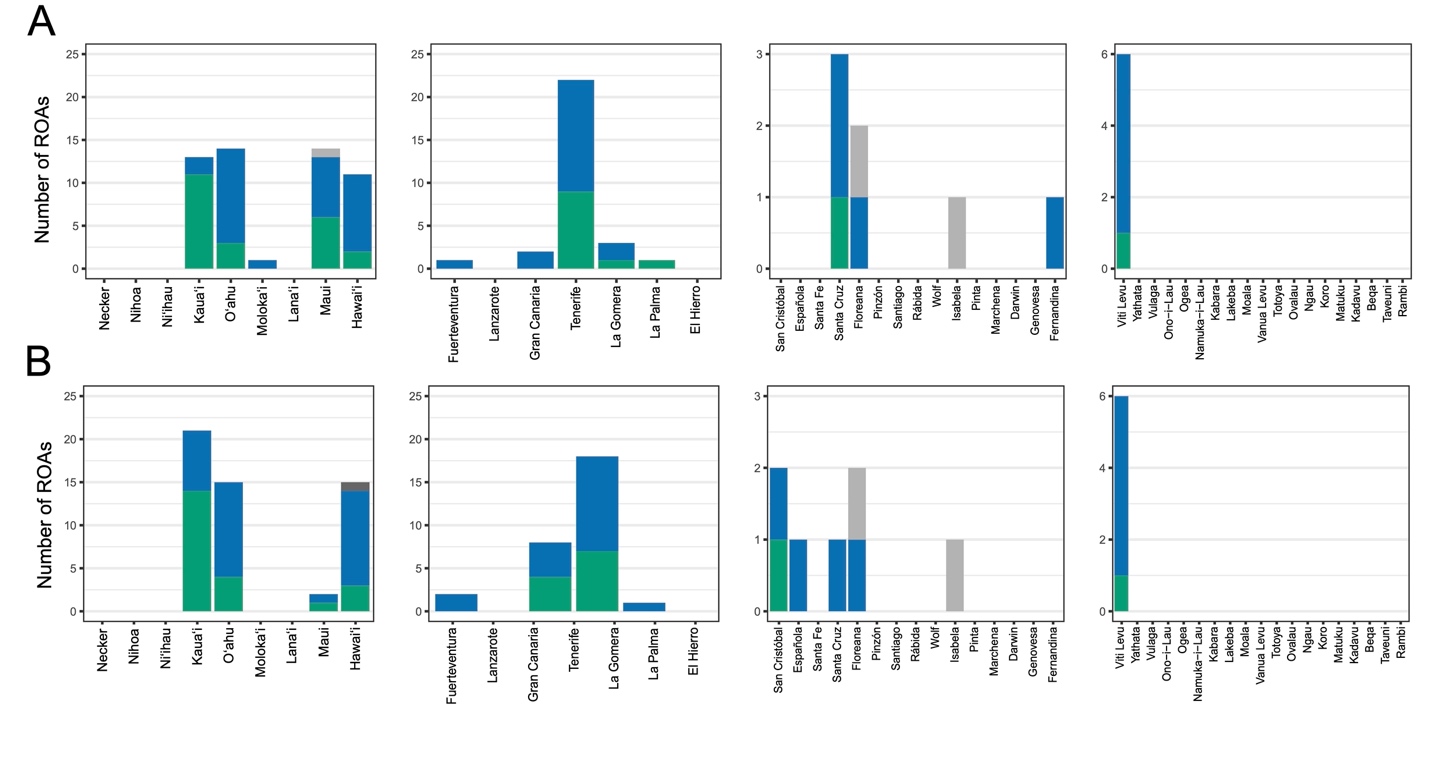

**Figure S4.** Number of ROAs per island that have their highest Island S_ROA_ (A) and highest number of Single Island Endemics (B) on the island. For instance, 22 ROAs reach their maximum Island S_ROA_ on Tenerife, and 18 ROAs reach their maximum number of SIE on this same island. Colors show the type of taxon (invertebrates=blue, plants=green, vertebrates=grey). We did not display the results for the Galápagos finches as six islands arrived first for highest Island S_ROA_ (for SIE, the maximum richness was reached in Genovesa).
